# GenVinci reconstructs 3D molecular tissue scenes from histology to map senescence across aging and disease

**DOI:** 10.64898/2026.11.17.767412

**Authors:** Xingjian Chen, Charles Comiter, Haochun Huang, Yinghan Wu, Ying-Li Su, Judy Zhou, Francesco Monticolo, Max Condouret, Styliani Sapantzi, Jianhuan Qi, Xuemeng Li, Yanwan Cao, Yanran Zhou, Ke Zhang, Ryota Tanaka, Kristina Todorova, Stefano Sol, Justin Ziegler, Denny Sakkas, Christine Lian, Shadmehr Demehri, Anna Mandinova, Wei Min, Jonathan C. Silverstein, Hemali Phatnani, Zhixun Dou, Jeon Woong Kang, Heng Yang, Aviv Regev, Collin M. Stultz, the NIH SenNet Consortium, Peter T. C. So, Jian Shu

## Abstract

Spatial omics has generated measurements at unprecedented scale, yet computational approaches remain largely confined to individual modalities and 2D representations. Here we assemble ImageOmicsNet-300M, a corpus of 300 million single-cell and spatial transcriptomic profiles, including 24.9 million H&E-predicted virtual spatial transcriptomics profiles, and 4,623 H&E sections. Of these H&E sections, we generated 1,782 from aging mouse organs and serially sectioned human skin spanning five decades, with matched spatial transcriptomics and snRNA-seq. Building on this resource, we develop GenVinci, a multimodal tissue foundation model that reconstructs 3D tissue scenes from routine 2D histology. GenVinci predicts spatial gene expression from H&E with zero-shot generalization and maps senescence-associated states across 85.8 million cells, revealing cell-type-specific patterns and disease-associated remodeling. Integration with human genetics implicates suprabasal keratinocytes in atopic dermatitis and keratinocyte carcinomas, linked to higher and lower p16 and p21 expression, respectively. These findings establish GenVinci as a framework mapping senescence in 3D across scales.

## INTRODUCTION

Cells do not act in isolation but exist within structured tissue scenes, in which molecular programs, cellular states, tissue morphology, and local microenvironments collectively shape biological function, development, aging, and disease^1,2^. Recent advances in spatial omics have enabled high-plex molecular profiling while preserving spatial context, generating increasingly rich multimodal measurements of tissue organization across molecular, cellular, and tissue scales^3–6^. However, reconstructing and interpreting coherent 3D tissue scenes, which are cellular-resolution representations that integrate individual cell states, their spatial organization, and surrounding tissue context in three dimensions across an entire tissue volume, extending beyond the local niche or microenvironment of any single cell, remains a fundamental challenge^5,7,8^.

Despite rapid advances in pathology and transcriptomics, existing approaches remain fragmented across modalities and scales. Spatial transcriptomic technologies provide rich molecular information but are costly, heterogeneous across platforms, and largely restricted to two-dimensional tissue sections, limiting their ability to characterize tissue organization at scale^9,10^. Conversely, routine histology preserves tissue architecture and is widely available across common clinical and research settings but lacks direct molecular measurements^11^. Consequently, the molecular and cellular architectures of tissues remain partially understood.

Computational methods have sought to bridge these modalities by learning transferable representations of cellular states and predicting molecular information from histological images^12–15^. Existing single-cell and spatial foundation models have performed well on classical transcriptomic tasks such as cell-type annotation and batch integration^16–19^. However, most are trained on dissociated single-cell RNA sequencing (scRNA-seq) data and lack explicit spatial or morphological grounding, limiting the biological contexts to which they transfer^16,17^. Recent histology-to-transcriptomics prediction approaches have shown promise but are typically trained on relatively small datasets; therefore, they struggle to generalize beyond the tissues or disease conditions on which they were developed^20–22^. These limitations have hindered the development of unified foundation models capable of learning broadly applicable representations for reconstructing and interrogating 3D tissue structure.

Here we present GenVinci, a multimodal tissue foundation model that reconstructs 3D tissue scenes from routine histology, built on ImageOmicsNet-300M, a curated image-transcriptomic corpus comprising more than 300 million cells and spatial profiles and 4,623 H&E sections, spanning two species, 82 tissue types, 55 transcriptomic technologies, and a range of biological conditions. Unlike existing transcriptomic foundation models that primarily treat cells as independent transcriptional entities^16,17^, GenVinci models cells as components of tissue scenes by incorporating modality, assay, organism, and tissue context directly into representation learning, and employs three complementary pretraining objectives to learn representations that capture intrinsic cell states, spatial organization, and tissue-level context. We further align the learned representations with morphological features to develop a generalizable H&E-to-transcriptome model, enabling *in silico* reconstruction of transcriptomic states, cellular identities, and spatial tissue organization directly from routine H&E images.

We evaluate GenVinci on held-out public and newly generated single-cell and spatial transcriptomic benchmarks spanning distinct tissues, platforms, and biological conditions, and apply it to 1,782 newly generated H&E sections in human and mouse aging to reconstruct virtual spatial transcriptomes comprising approximately 25 million cells. Across established representation-learning benchmarks, GenVinci outperforms existing transcriptomic foundation models (Geneformer, scGPT, Nicheformer, and scGPT-spatial^16–19^) in cell-type annotation, spatial domain identification, and batch integration, while further improving age prediction relative to SpatialAgingClock^23^ and senescence identification compared with single-marker^24^, gene-set-based^25,26^, and machine-learning-based approaches^27,28^. We then leverage the learned representations for biological discovery by mapping senescence-associated states across 85.8 million cells in ImageOmicsNet-300M and build SenNet-100, an AI-derived single-cell signature of senescence that separates a canonical stress response^29^ from a cytoskeletal–adhesion program. By transferring GenVinci representations to histology through representation alignment (REPA)^30^, we further reconstruct spatial transcriptomic states from H&E and extend these predictions across hundreds of serial human skin sections across ages to generate cellular-resolution 3D tissue scenes. Integration of these reconstructed tissue scenes with human genetic data across five skin diseases further enables spatial mapping of disease-associated genetic risk^31^, localizing the risk for inflammatory diseases and keratinocyte carcinomas to suprabasal keratinocytes across donors and ages, and providing a framework for interrogating aging and disease across molecular, cellular, and tissue scales.

Together, GenVinci establishes a general foundation for reconstructing and interrogating 3D tissue scenes, suggesting that senescence-associated programs of aging and disease are organized by cell identity and tissue context rather than as a uniform cellular state.

## RESULTS

### Construction of ImageOmicsNet-300M and the GenVinci foundation model

GenVinci is a foundation model built on ImageOmicsNet-300M, a large-scale curated multimodal image-transcriptomic corpus assembled from more than 20 public resources, consortium-wide data from the NIH SenNet Consortium, and newly generated datasets, together spanning more than 2,000 studies and publications^32–36^ (Figure 1A, Figure S1A, and Table S1). The corpus comprises 300.2 million transcriptomic profiles, including 275.3 million experimentally measured scRNA-seq and spatial profiles (58.7 million from the SenNet Consortium from 517 donors in 1,150 datasets contributed by 11 Tissue Mapping Centers and 4 Technology Development and Application teams: 3.7 million dissociated and 55.1 million spatial) and 24.9 million H&E-predicted virtual spatial profiles, together with 4,623 H&E sections (2,841 publicly or consortium collected and 1,782 newly generated). It spans two species, 82 tissue types, 33 scRNA-seq assays, 22 spatial transcriptomic technologies, and 1,420 cell types. We also newly generated STARmap data from mouse tongue (two samples, one young and one old; 4,858 cells), kidney (two samples, one young and one old; 30,157 cells), placenta (one E14.5 sample; 10,833 cells), and human skin (four sections, three from a 23-year-old donor and one from a 71-year-old donor; 21,108 cells), together with a human skin snRNA-seq dataset (24,153 nuclei) (Methods). The scale and diversity of ImageOmicsNet-300M provide broad biological coverage, including aging- and senescence-associated states from the SenNet Consortium^37^ and our newly generated mouse and human skin aging atlases, supporting generalizable tissue scene representation learning across diverse biological contexts.

**Figure 1.**
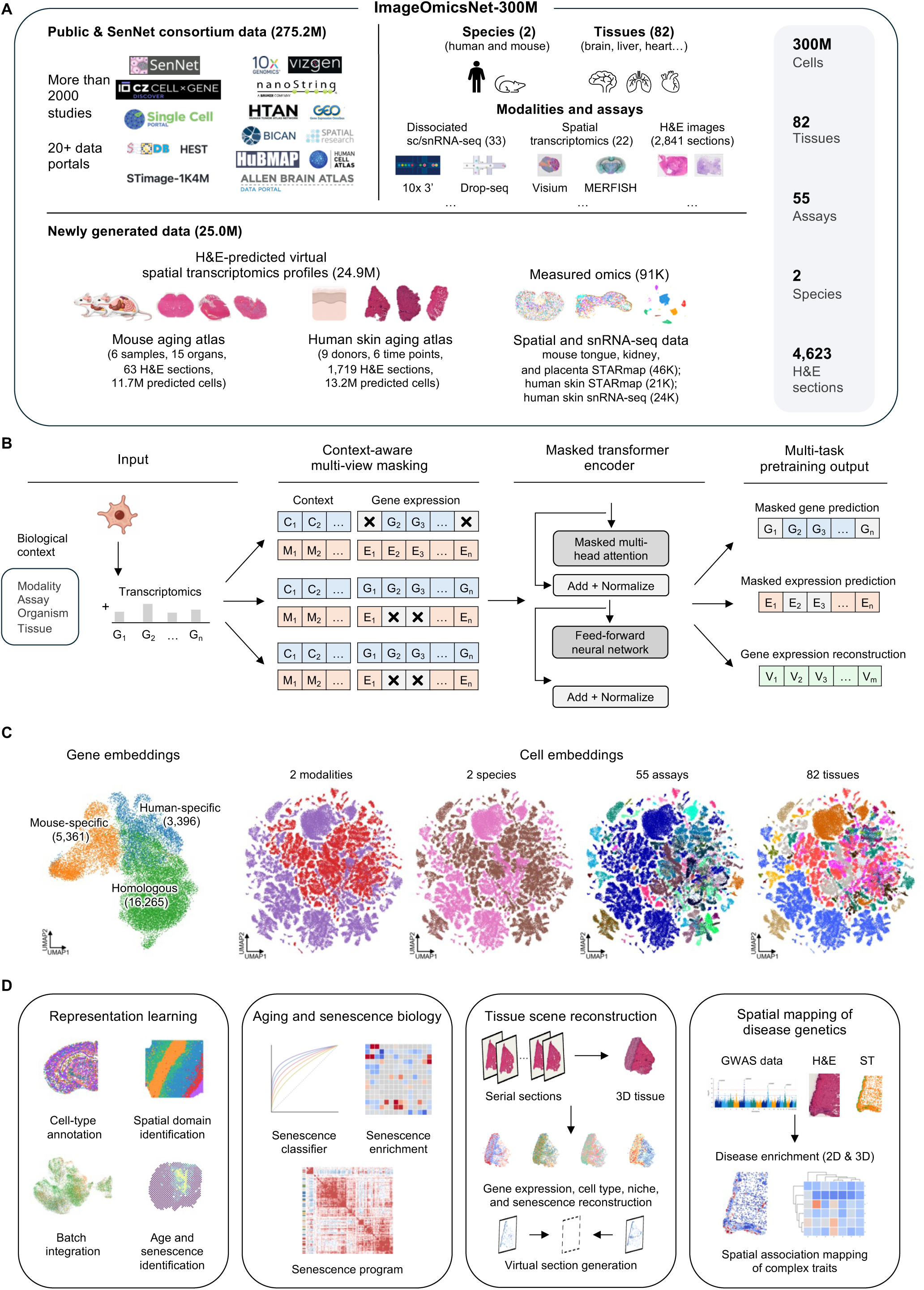
ImageOmicsNet-300M and GenVinci enable transferable tissue scene representations. **A,** Overview of the ImageOmicsNet-300M corpus, integrating public, consortium (SenNet), and newly generated datasets across two species, 82 tissues, and 55 transcriptomic technologies. The corpus comprises 300.2 million scRNA-seq and spatial profiles (275.3 million experimentally measured and 24.9 million H&E-predicted virtual spatial profiles) and 4,623 H&E sections. **B,** Overview of the GenVinci pretraining framework. Transcriptomic profiles and biological context, including modality, assay, organism, and tissue, are jointly encoded through context-aware multi-view masking and processed by a masked transformer encoder. The model is trained using three self-supervised objectives: masked gene prediction, masked expression prediction, and gene expression reconstruction. **C,** Representations learned during GenVinci pretraining. Left, UMAP of gene embeddings, showing homologous, human-specific, and mouse-specific genes. Right, UMAPs of cellular representations from a random subsample of 100,000 cells, colored by modality, species, assay, and tissue. **D,** Overview of downstream applications enabled by ImageOmicsNet-300M and GenVinci: representation learning, aging and senescence biology, tissue scene reconstruction, and spatial mapping of disease genetics.

We first pretrained GenVinci on 211.0 million experimentally measured scRNA-seq and spatial profiles from ImageOmicsNet-300M, prior to incorporation of the expanded SenNet data release, which was not used for model training or analyses in this study (Figure S1A and Methods). The 6.3 million spatial profiles with paired H&E images (from HEST-1k^32^ and STimage-1K4M^36^) were reserved for the histology-to-transcriptome analysis, and all datasets newly generated or used for downstream evaluation were manually excluded from pretraining. Instead of treating cells as independent transcriptional entities, GenVinci explicitly models cells as components of larger tissue scenes defined by both contextual and molecular relationships (Figure 1B and Methods). To capture these dependencies, we incorporated modality, assay, organism, and tissue information into the encoder, allowing contextual information to interact directly with gene expression during pretraining (Figure 1B and Figure S1B). GenVinci employs a context-aware multi-view masking strategy^38^ in which different views of each cell are generated by masking either gene identities or expression levels while preserving shared contextual information (Figure 1B). The model is optimized using three complementary objectives—masked gene prediction^17^, masked expression prediction^16^, and gene expression reconstruction (Methods)—which respectively capture gene co-expression context, fine-grained expression differences, and information lost to sequence truncation and expression binning.

To examine whether biologically meaningful representations were learned during pretraining, we visualized the learned gene and cell embeddings using uniform manifold approximation and projection (UMAP)^39^ (Figure 1C and Figure S1D). Gene embeddings reflected species-level organization consistent with orthology, with conserved homologous genes positioned between partially separated clusters of human- and mouse-specific genes. Beyond this species-level organization, the learned representations further captured functional relationships among genes. Canonical gene families, including ribosomal, collagen, keratin, chemokine, histone, and mitochondrial genes, formed coherent local neighborhoods in the embedding space and exhibited elevated within-family cosine similarity (Figure S1C). Cell embeddings were well integrated across modalities, species, assays, and tissues, while preserving distinct clusters corresponding to major cell populations, including immune, neuronal, epithelial, and stromal cells (Figure 1C and Figure S1D). These results indicate that GenVinci organizes genes and cells into biologically coherent representation spaces that preserve both evolutionary and cellular relationships and provide a foundation for downstream analyses of tissue organization and cellular states.

We also summarize the range of analyses enabled by GenVinci in an overview schematic (Figure 1D). These applications span four interconnected domains: representation learning, aging and senescence biology, tissue scene reconstruction, and spatial mapping of disease genetics. Together, these applications position GenVinci as a unified framework for reconstructing and interrogating tissue scenes across molecular, cellular, and tissue scales.

### GenVinci learns transferable representations of cellular and spatial tissue organization

To evaluate the quality and transferability of the learned representations, we first benchmarked GenVinci against existing single-cell and spatial foundation models (Geneformer^17^, scGPT^16^, Nicheformer^19^, and scGPT-spatial^18^) across three widely used downstream tasks: cell-type annotation, spatial domain identification, and batch integration (Figure 2 and Methods). These tasks evaluate complementary aspects of representation quality, including supervised biological classification, recovery of spatial tissue organization, and robustness to technical variation.

**Figure 2.**
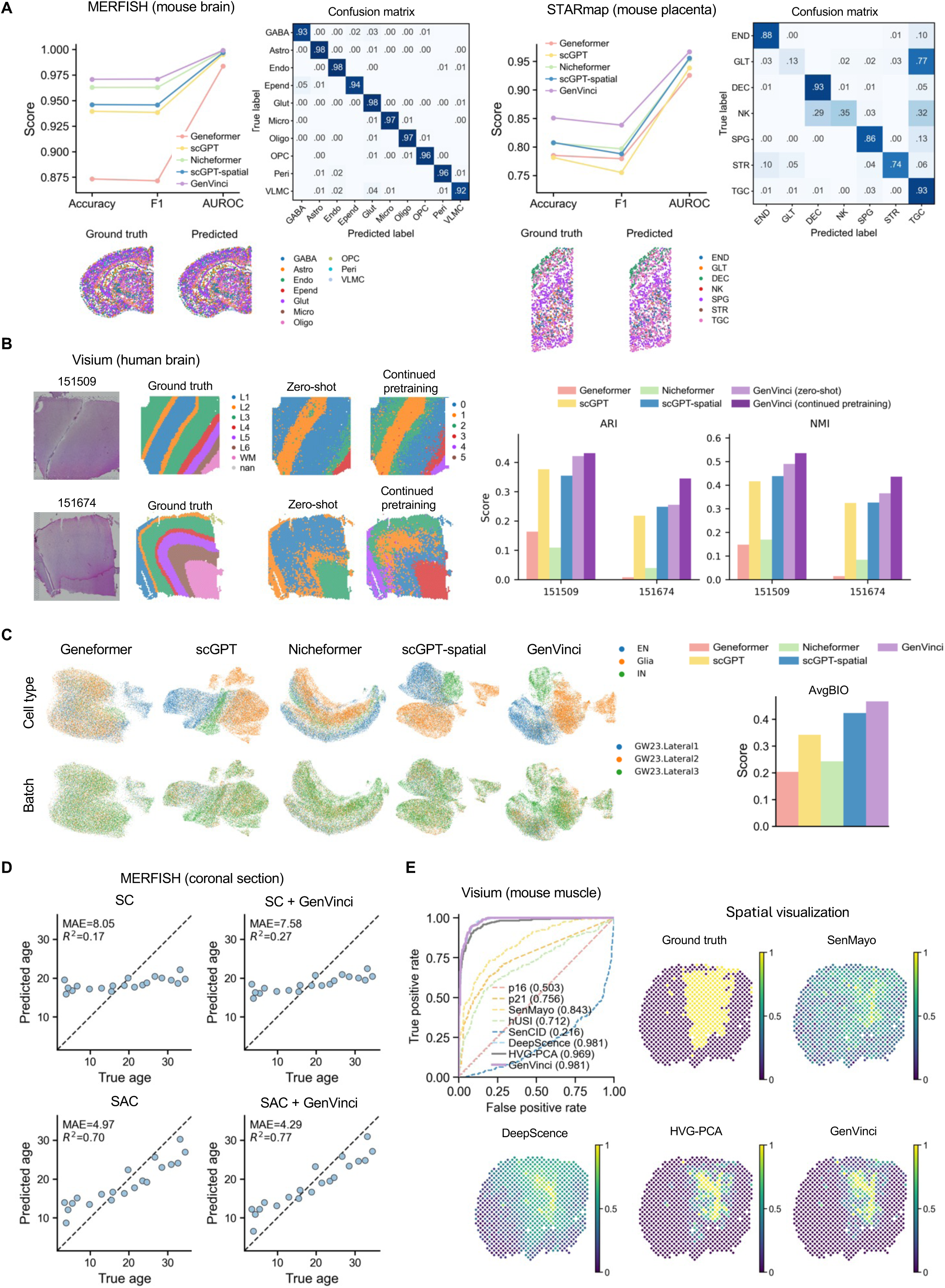
GenVinci learns transferable representations for cell identity, spatial organization, aging, and senescence. **A,** Cell-type annotation on MERFISH mouse brain and STARmap mouse placenta datasets. For each dataset: left, quantitative comparison of accuracy, F1 score, and AUROC across Geneformer, scGPT, Nicheformer, scGPT-spatial, and GenVinci; right, GenVinci confusion matrix (values are proportions of each true label); bottom, ground-truth and GenVinci-predicted cell-type maps. Cell-type abbreviations—MERFISH: GABA, GABAergic neuron; Astro, astrocyte; Endo, endothelial cell; Epend, ependymal cell; Glut, glutamatergic neuron; Micro, microglial cell; Oligo, oligodendrocyte; OPC, oligodendrocyte precursor cell; Peri, pericyte; VLMC, vascular leptomeningeal cell. STARmap: END, endothelial cell; GLT, glandular trophoblast; DEC, maternal decidua; NK, NK cell; SPG, spongiotrophoblast; STR, stromal cell; TGC, trophoblast giant cell. **B,** Spatial domain identification on two human Visium brain samples (151509 and 151674). Left, H&E images and manually annotated cortical layers (L1–L6 and white matter, WM; nan, unannotated spots). Middle, zero-shot clustering from GenVinci. Right, clustering after continued pretraining. Predicted clusters (0–5) are unsupervised and are not label-matched to annotated layers. Bar plots summarize ARI and NMI across both sections for all methods. **C,** Batch integration of the MERFISH human developmental thalamus dataset across Geneformer, scGPT, Nicheformer, scGPT-spatial, and GenVinci. UMAP visualizations are colored by cell type (top: EN, excitatory neurons; IN, interneurons; Glia, glial cells) or batch (bottom: three lateral sections from a single GW23 donor). AvgBIO quantifies preservation of biological variation after integration. **D,** Age prediction on the MERFISH coronal mouse brain dataset using normalized single-cell expression (SC), SpatialAgingClock (SAC), and the corresponding GenVinci-based variants (SC + GenVinci, SAC + GenVinci). Scatter plots show mouse-level predicted versus ground-truth age, with MAE and *R*² indicated. **E,** Senescence identification in the mouse muscle Visium dataset. Left, ROC curves across competing methods (dashed lines, marker-based (p16 and p21), gene-set scoring (SenMayo and hUSI), and machine-learning-based (SenCID and DeepScence) methods; solid lines, HVG-PCA and GenVinci); AUROC values are indicated. Right, spatial distributions of predicted senescence scores for each approach alongside ground-truth annotations. ARI, adjusted Rand index; NMI, normalized mutual information; MAE, mean absolute error; AUROC, area under the receiver operating characteristic (ROC) curve; HVG, highly variable gene; GW23, gestational week 23.

For cell-type annotation, we evaluated GenVinci on a public mouse brain multiplexed error-robust fluorescence in situ hybridization (MERFISH) dataset^40^ and our in-house mouse placenta spatially resolved transcript amplicon readout mapping (STARmap)^5^ dataset, both held out from pretraining and representing distinct tissues, spatial technologies, and gene panels (Figure 2A). For a fair comparison, all methods were evaluated using the same linear probing framework (Methods). On the MERFISH dataset, GenVinci achieved an AUROC of 0.999 and both an accuracy and F1 score of 0.971, surpassing the strongest baseline (Nicheformer; AUROC 0.998, accuracy 0.963, F1 0.963; Figure 2A). On the STARmap dataset, GenVinci achieved larger gains, improving accuracy from 0.808 (scGPT-spatial) to 0.851, F1 score from 0.797 (Nicheformer) to 0.838, and AUROC from 0.956 (scGPT-spatial) to 0.967 (Figure 2A). Predicted spatial distributions closely recapitulated the ground-truth tissue organization, and confusion-matrix analyses revealed improved discrimination across major cellular populations relative to other models (Figure 2A and Figure S2A). These findings indicate that GenVinci preserves fine-grained cellular identity while generalizing across distinct spatial transcriptomic platforms and tissue contexts.

For spatial domain identification, we used two human dorsolateral prefrontal cortex samples with manually annotated cortical layers and white matter regions^41^. To specifically evaluate whether pretrained representations encode tissue-domain information, spatial-coordinate information was excluded from all methods, and clustering was performed solely on the learned representations (Methods). Across both samples, GenVinci achieved the best recovery of spatial domains (Figure 2B), obtaining adjusted Rand index (ARI)/normalized mutual information (NMI) scores of 0.42/0.49 and 0.26/0.36 for samples 151509 and 151674, respectively. We further assessed the benefit of task-specific adaptation through continued pretraining on the target dataset without using any annotation labels, compared with zero-shot representations (Methods). This unsupervised continued pretraining further improved ARI/NMI scores to 0.43/0.53 and 0.35/0.44 for samples 151509 and 151674, respectively, yielding more coherent spatial domains and sharper boundaries between neighboring cortical layers (Figure 2B). Visual inspection confirmed that GenVinci embeddings showed greater correspondence with annotated cortical layers and white matter regions than those of the other methods (Figure 2B and Figure S2B). These results suggest that GenVinci captures molecular programs intrinsically linked to tissue organization, even without explicit spatial-coordinate information.

We next assessed batch integration and benchmarked GenVinci on a MERFISH human developmental thalamus dataset comprising three batches and three annotated cell types^42^ (Methods). Across all methods, GenVinci achieved the highest overall average biological conservation score (AvgBIO = 0.47), outperforming Geneformer (0.20), scGPT (0.34), Nicheformer (0.24), and scGPT-spatial (0.42) (Figure 2C, right). This advantage was reflected across multiple metrics, including cluster-label agreement (NMI = 0.43, ARI = 0.42) and average silhouette width (ASW) for cell-type label (ASW_label = 0.54) (Figure S2C). UMAP visualizations showed that GenVinci separated major biological populations more clearly, such as excitatory neurons (EN), interneurons (IN), and glial cells (Glia), while maintaining effective mixing across batches (Figure 2C, left). Importantly, batch-conservation metrics, including ASW batch and graph connectivity, remained comparable across all methods (Figure S2C, bottom), indicating that the improved biological conservation achieved by GenVinci was not driven by overcorrection or loss of batch alignment.

Moreover, given the extensive aging- and senescence-associated resource in ImageOmicsNet-300M, we asked whether GenVinci captures these biological signals without task-specific training. We evaluated two tasks rarely examined for transcriptomic foundation models: age prediction on two MERFISH mouse brain datasets^23^ and senescence identification on two spatial transcriptomic datasets^43,44^ (Figure 2D,E; Figure S2D,E; Figure S3; Methods).

For age prediction, on the coronal dataset, we trained a separate model for each cell type using leave-one-mouse-out cross-validation, holding out all cells from one mouse for testing in each fold (Figure S3A; Methods). To test cross-dataset generalization, we also trained on the coronal dataset and tested on an independent sagittal dataset from a different cohort. We compared four inputs: normalized single-cell expression (SC), SpatialAgingClock (SAC)^23^, which smooths SC across spatial neighbors, and their variants in which SC is replaced with GenVinci embeddings (SC + GenVinci and SAC + GenVinci). All comparisons used the same cross-validation folds and regressor. In the SC setting, GenVinci reduced the mean absolute error (MAE) from 8.05 to 7.58 and increased *R*² from 0.17 to 0.27 on the coronal dataset (Figure 2D, top), with similar improvements on the independent sagittal dataset (MAE, 6.85 to 6.47; *R*², 0.18 to 0.32; Figure S2D, top). In the SAC setting, GenVinci similarly improved performance on both the coronal dataset (MAE, 4.97 to 4.29; *R*², 0.70 to 0.77; Figure 2D, bottom) and the sagittal dataset (MAE, 3.53 to 3.24; *R*², 0.76 to 0.79; Figure S2D, bottom). GenVinci also reduced prediction error for most cell types in both settings and both datasets (Figure S3B). Importantly, because GenVinci improved performance both with and without spatial smoothing, these gains are attributable to the learned representation rather than to spatial context alone.

For senescence identification, senescent cells were labeled following the DeepScence protocol^27^. We benchmarked GenVinci against marker-based methods (*CDKN2A*/p16 and *CDKN1A*/p21)^24^, gene-set-based methods (SenMayo^26^ and hUSI^25^), and machine-learning-based methods (SenCID^28^ and DeepScence^27^). As a matched supervised baseline, we also trained a logistic regression classifier on principal components of highly variable genes (HVG-PCA), using the same labels and folds (Methods). GenVinci outperformed the matched supervised baseline on both spatial datasets (AUROC: mouse muscle^43^, 0.981 vs. 0.969 for HVG-PCA; human brain^44^, 0.956 vs. 0.634), with the larger margin observed on the Stereo-seq human brain dataset (Figure 2E and Figure S2F). Spatial visualization further showed that GenVinci scores localized to the annotated senescent regions, whereas marker-based and gene-set scores showed limited spatial correspondence (Figure 2E, Figure S2E, and Figure S3C).

Collectively, these benchmarks show that GenVinci learns biologically grounded transcriptomic representations that generalize across spatial technologies, tissues, and downstream tasks, while preserving signals associated with aging and cellular senescence. By capturing both classical transcriptomic properties and aging-related states, GenVinci provides a foundation for downstream biological discovery, molecular reconstruction, and tissue scene analysis.

### GenVinci reveals context-dependent senescence remodeling and a transferable senescence module

Leveraging the extensive representation of healthy, aging, and disease tissues in ImageOmicsNet-300M, we then asked whether the learned GenVinci representation could be used to systematically characterize cellular senescence at atlas scale. To this end, we developed a weakly supervised framework for large-scale senescence mapping across ImageOmicsNet-300M (Figure 3A). We first defined high-confidence positive and negative seed labels within strata jointly defined by species, cell type, donor unit, and within-stratum sequencing-depth decile, using expression of the canonical senescence markers *CDKN2A* (p16) and *CDKN1A* (p21)^24^. Seed-labeled cells were pooled across strata to train a supervised classifier in the shared GenVinci representation space, with the *CDKN1A* and *CDKN2A* tokens masked before encoding to prevent label leakage. Applying this classifier to the 85.8 million cells in ImageOmicsNet-300M with curated cell-type and disease annotations (Figure S4A), spanning 2 species, 70 organs, 930 cell types, and 141 diseases, generated a unified continuous senescence score. For downstream analyses, we designated the 5% of cells with the highest predicted senescence scores as senescent, which serves as an operational threshold rather than an estimate of the true senescent fraction (Methods). Likewise, the two canonical markers served only as anchors for learning the senescence representation, not as a definition of the senescent state.

**Figure 3.**
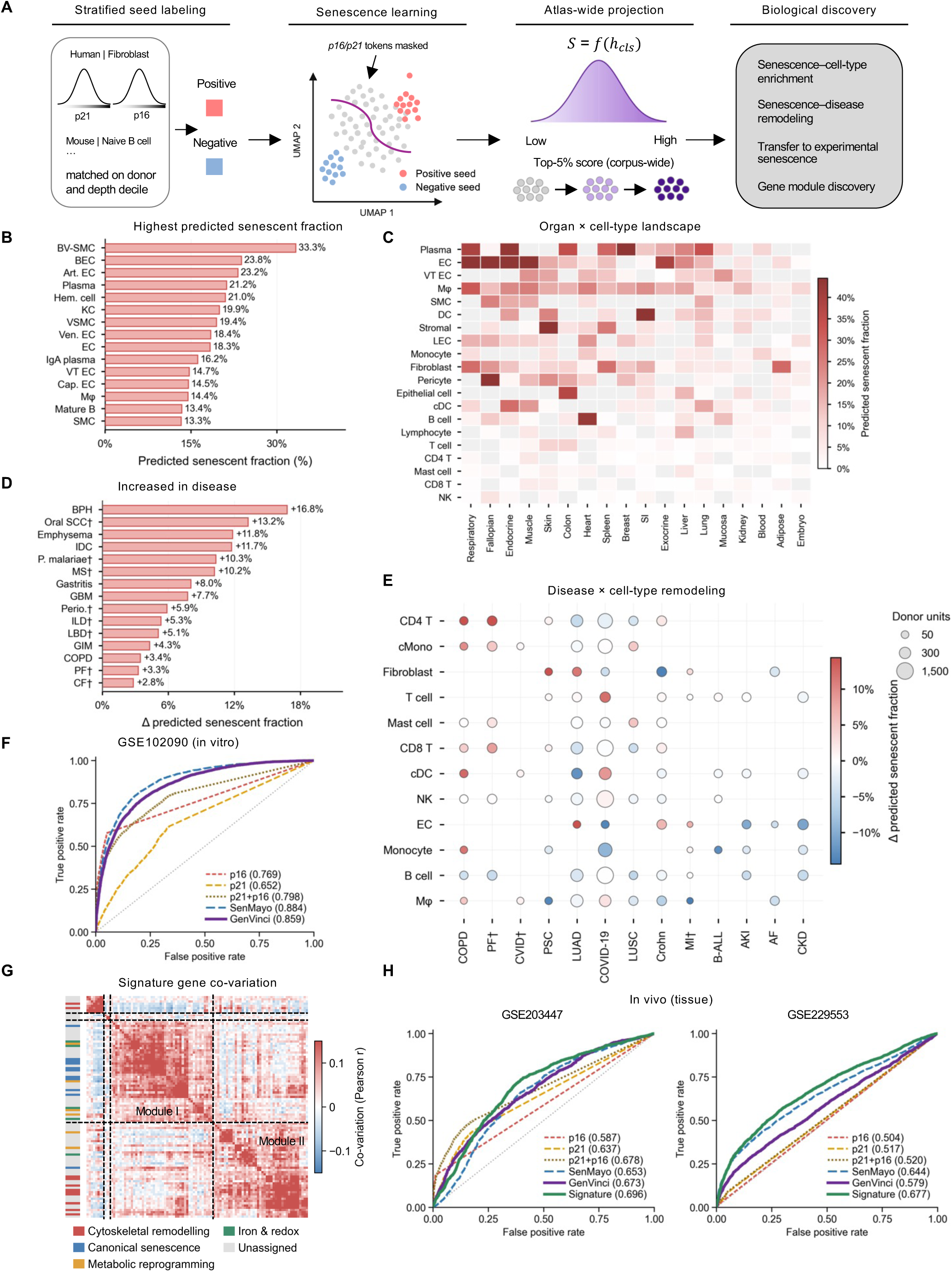
A weakly supervised classifier charts the predicted senescent fraction across an 85.8-million-cell atlas and transfers to experimentally induced senescence. **A,** Overview of the senescence analysis framework. Left, stratified seed labeling using *CDKN1A* (p21) and *CDKN2A* (p16) detection as weak labels. Middle left, a classifier is trained on CLS embeddings computed with the p16 and p21 tokens removed. Middle right, the learned score *S* is projected across the corpus and the top 5% of cells are retained as predicted senescent. Right, the four downstream analyses enabled by the projection. **B,** The 15 cell types with the highest predicted senescent fraction (proportion of cells in the top 5% of S) in healthy samples, among the 86 cell types passing quality control. Each observation is a cell type; bars show donor-weighted means across groups. **C,** Organ × cell-type landscape of predicted senescent fraction in healthy samples, for the 20 cell types spanning the most organs across the 18 organs containing them. Rows are ordered by overall predicted senescent fraction and columns by mean predicted senescent fraction; light gray cells indicate combinations with no qualifying group. **D,** The 15 diseases with the largest increase in predicted senescent fraction relative to matched healthy samples. Daggers denote diseases for which the estimate derives from a single dataset. **E,** Disease × cell-type remodeling for the 13 diseases and 12 cell types with the broadest mutual coverage. Point color indicates the donor-weighted change in predicted senescent fraction relative to matched healthy samples and point size the total number of donor units contributing to each estimate; daggers as in D. **F,** ROC curves for senescent-cell identification in the GSE102090 *in vitro* dataset (S vs. NS; S, senescent; NS, non-senescent) by marker-based detection (p16, p21, and their union), gene-set scoring (SenMayo), and the senescence classifier (GenVinci); AUROC values are indicated. **G,** Co-variation structure of SenNet-100 (hierarchically clustered Pearson correlations). Black lines delimit the four clusters (*k* = 4); the left color bar shows an independent manual functional annotation not used for clustering. Clusters 3 (46 genes; stress–inflammatory) and 4 (43 genes; cytoskeletal–adhesion) define Module I and Module II. **H,** ROC curves for two *in vivo* tissue datasets (GSE203447 and GSE229553), comparing the same approaches as in F together with SenNet-100 (labeled “Signature”). Cell-type abbreviations—BV-SMC, blood vessel smooth muscle cell; BEC, blood vessel endothelial cell; Art. EC, endothelial cell of artery; Ven. EC, vein endothelial cell; Cap. EC, capillary endothelial cell; VT EC, endothelial cell of vascular tree; LEC, endothelial cell of lymphatic vessel; VSMC, vascular associated smooth muscle cell; SMC, smooth muscle cell; EC, endothelial cell; KC, keratinocyte; Mφ, macrophage; Plasma, plasma cell; Hem. cell, hematopoietic cell; cDC, conventional dendritic cell; cMono, classical monocyte. Disease abbreviations—BPH, benign prostatic hyperplasia; Oral SCC, oral cavity squamous cell carcinoma; IDC, invasive ductal breast carcinoma; *P. malariae*, *Plasmodium malariae* malaria; MS, multiple sclerosis; GBM, glioblastoma; Perio., periodontitis; ILD, interstitial lung disease; LBD, Lewy body dementia; GIM, gastric intestinal metaplasia; COPD, chronic obstructive pulmonary disease; PF, pulmonary fibrosis; CF, cystic fibrosis; CVID, common variable immunodeficiency; PSC, primary sclerosing cholangitis; LUAD, lung adenocarcinoma; LUSC, squamous cell lung carcinoma; Crohn, Crohn disease; MI, myocardial infarction; B-ALL, B-cell acute lymphoblastic leukemia; AKI, acute kidney injury; AF, atrial fibrillation; CKD, chronic kidney disease.

We then established a physiological reference landscape of senescence across healthy tissues, aggregating 2,135 organ–cell-type groups (≥1,000 cells each; 61 organs) into 722 cell types. Predicted senescent fractions across cell types were strongly right-skewed (median 2.9%), with most showing low baseline levels and a smaller subset markedly elevated (Figure S4B). Ranking sufficiently represented cell types (Methods) revealed substantial cell-type-associated variation in predicted senescent fractions (Figure 3B). Vascular populations dominated the high-baseline end of the landscape, with blood vessel smooth muscle cells (BV-SMC; 33.3%), blood vessel endothelial cells (BEC; 23.8%), and arterial endothelial cells (Art. EC; 23.2%) ranking highest, together with plasma cells (21.2%), hematopoietic cells (Hem. cell; 21.0%), and keratinocytes (KC; 19.9%), whereas naive and resting lymphoid populations occupied the low end, including naive CD8⁺ T cells (0.05%), CD56dim NK cells (0.30%), and naive CD4⁺ T cells (0.70%; Figure S4C). Notably, this vascular enrichment emerged without any lineage-specific input to the model and is consistent with previous reports of senescent cell accumulation in the vasculature^45,46^.

We next examined how the baseline states varied across tissue contexts. Cross-organ analysis revealed that the cell-type ordering was largely preserved across organs (Figure 3C). Endothelial and smooth muscle populations showed elevated baseline senescence in respiratory, endocrine, muscle, skin, and cardiac tissues alike, whereas T, CD8⁺ T, and NK populations remained near the floor of the scale in every organ examined. Stromal and fibroblast populations displayed the greatest organ-dependent variation, consistent with their tissue-specialized identities^47^. These patterns suggest that cell-type identity is the dominant determinant of baseline senescence in healthy tissue, with tissue context acting as a secondary modifier rather than an independent axis.

Having established the physiological landscape, we next investigated senescence remodeling across disease states, computed within matched species–organ–cell-type strata. Disease effects were bidirectional and centered slightly below zero across 77 diseases (median −1.5%; Figure S4D), indicating that pathological states do not uniformly elevate the predicted senescent fraction. Benign prostatic hyperplasia (BPH; +16.8%), oral cavity squamous cell carcinoma (Oral SCC; +13.2%), pulmonary emphysema (+11.8%), and breast invasive ductal carcinoma (IDC; +11.7%) ranked among the highest-scoring disease contexts (Figure 3D), whereas malignant ovarian serous tumor (OSC; −28.0%), endocrine pancreas disorder (−11.6%), and atherosclerosis (−9.4%) ranked among the lowest (Figure S4E). The negative tail was enriched for proliferative malignancies, in which expansion of highly proliferative compartments is expected to dilute an arrest-associated phenotype. Rather than exhibiting uniform activation, disease-associated senescence proved highly cell-type specific and frequently discordant in direction within a single disease (Figure 3E). In COVID-19, predicted senescent fractions were reduced in CD4⁺ T cells, classical monocytes, and B cells but elevated in fibroblasts and conventional dendritic cells. Such opposing shifts can offset one another in disease-level summaries, obscuring substantial cell-type-specific remodeling.

We next asked whether the learned classifier generalizes to senescence states beyond those defined by marker detection. To this end, we evaluated it on four comparisons of experimentally induced senescence from two external *in vitro* datasets^48,49^ (Methods). The predicted senescence score achieved a mean AUROC of 0.824 across the four comparisons (Figure 3F and Figure S4F–H). It performed comparably to SenMayo under spontaneous and replicative senescence (GSE102090, 0.859 vs. 0.884; GSE226225 replicative, 0.865 vs. 0.857) and substantially outperformed it under acute induction, where SenMayo performance dropped markedly (GSE226225 irradiation, 0.774 vs. 0.589; etoposide, 0.798 vs. 0.648). Notably, SenMayo is a literature-curated set of 125 senescence-associated genes^26^, whereas our classifier was supervised only by detection of two genes, both masked from the input before encoding. The predicted score further exceeded direct scoring of its own supervisory signal, with combined p16/p21 detection reaching a mean AUROC of 0.754 and only 0.659 under replicative senescence. These results indicate that, although initialized with p16/p21-based seed labels, the learned representation generalizes to experimentally induced senescence states beyond these marker genes.

We next quantified gene-wise associations between expression and predicted senescence across all species–cell-type strata and retained genes showing positive associations that were both widespread across strata and strong in magnitude, yielding a senescence signature comprising 100 genes (SenNet-100; Methods and Table S2). Hierarchical clustering of the within-stratum expression differences between high- and low-scoring cells partitioned SenNet-100 into two principal clusters (modules I and II) of 46 and 43 genes, together with two minor clusters of 8 and 3 genes (Figure 3G). Although no functional annotation was provided during clustering, 10 of 11 manually annotated canonical senescence genes segregated into the first module and 9 of 11 cytoskeletal genes into the second; the two remaining cytoskeletal genes, *DIAPH3* and *TUBB5*, instead joined a small proliferation-associated cluster, consistent with their mitotic rather than structural roles. Over-representation analysis showed that module I enriched for the canonical senescence-associated stress response (TNFα/NF-κB signaling, FDR = 8.8 × 10⁻³³; apoptosis, FDR = 2.2 × 10⁻¹⁵; hypoxia and interferon-γ responses; Figure S4J) and module II for cytoskeletal and adhesion programs (cytoskeleton, FDR = 1.2 × 10⁻⁷; focal adhesion, FDR = 4.4 × 10⁻⁷; Figure S4K). Module I additionally included 28 genes and module II 30 genes that had resisted manual assignment, indicating that GenVinci identifies coordinated gene programs beyond canonical marker genes, and that a cytoskeletal–adhesion program is separable from the secretory stress program with which senescence is conventionally identified.

Finally, we evaluated the classifier and SenNet-100 *in vivo*, where senescent cells are rare and transcriptionally overlap with inflammatory and stress states^29^. All methods fell within a narrow range in this setting (Figure 3H). Across two independent tissue datasets^50,51^, SenNet-100 achieved the highest AUROC, outperforming SenMayo (0.696 vs. 0.653 and 0.677 vs. 0.644), combined p16/p21 detection (0.678 and 0.520), and every individual marker, whereas the classifier performed comparably to or below SenMayo (0.673 and 0.579). This contrast likely reflects how each readout was derived. The classifier was trained to discriminate cells within matched strata, a structure closely mirrored by *in vitro* comparisons of treated and control cells but not by tissue, where donor, activation state, and microenvironment vary jointly. In contrast, SenNet-100 aggregates expression differences across many matched strata, making it less sensitive to such heterogeneity; however, it ranked acutely induced senescent cells below controls *in vitro* (AUROC 0.319 for irradiation and 0.218 for etoposide; Figure S4I). Together, these results suggest that the classifier and SenNet-100 are complementary, with the classifier better suited to controlled *in vitro* settings and SenNet-100 providing a more robust readout of senescence *in vivo*.

### Transfer of GenVinci representations to histology enables virtual spatial transcriptomics

Having shown that GenVinci captures biologically interpretable transcriptomic organization, we next asked whether these representations could be transferred to histology for virtual spatial transcriptomics. Histological images provide rich morphological information but lack direct molecular readouts. Existing approaches typically learn image features while directly regressing gene expression, without explicitly leveraging the structure of transcriptomic representations^13,21^. Such direct prediction can be sensitive to biological and technical distribution shifts across datasets, limiting out-of-distribution generalization.

To transfer the biological representations learned by GenVinci into tissue morphology, we developed GenVinci-2DST, a representation alignment framework built on 6.3 million paired histology–transcriptome profiles from ImageOmicsNet-300M (Figure S5A). Instead of directly predicting gene expression from H&E images, GenVinci first encodes spatial transcriptomics into robust representations that serve as biological supervision, while H&E image features extracted by a pathology foundation model (H-optimus-1^52^) are projected into the same latent space through an image encoder (Figure 4A). A decoder subsequently reconstructs gene expression from the aligned latent representation. This design utilizes GenVinci representations as an intermediate molecular supervision space between histology and gene expression reconstruction, with spatial information incorporated through a Spatial Refinement Module (SRM; Figure S5B and Methods).

**Figure 4.**
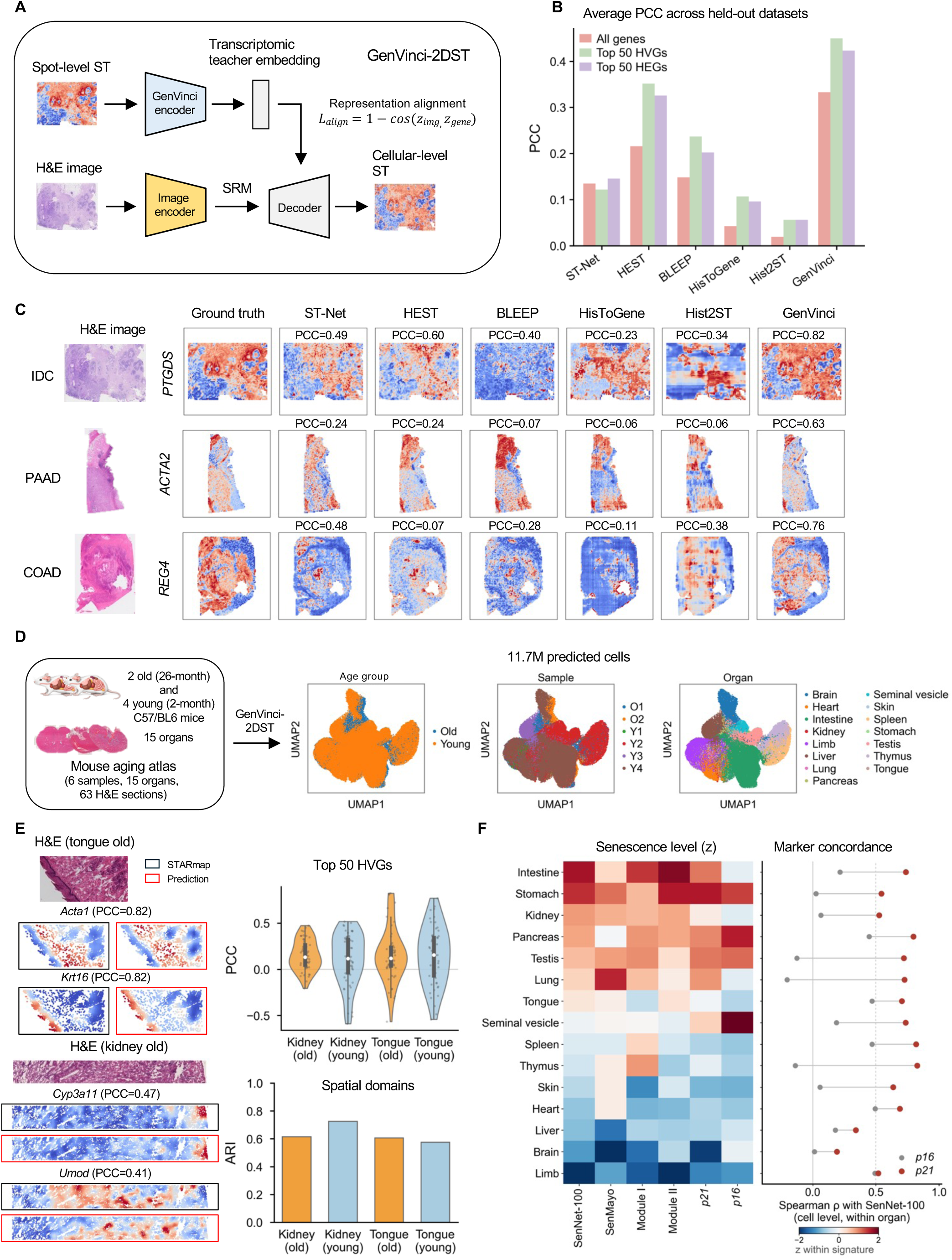
Transfer of GenVinci representations to histology enables virtual spatial transcriptomics and multi-organ molecular reconstruction. **A,** Schematic of GenVinci-2DST, in which GenVinci-derived transcriptomic teacher embeddings supervise the alignment of histology representations extracted by a pathology foundation model, followed by decoding into virtual cellular-level spatial transcriptomics. B, Comparison of gene-expression prediction performance between GenVinci-2DST and competing methods across held-out cancer datasets. Bars show mean Pearson correlation coefficients (PCC) for all genes, the top 50 highly variable genes (HVGs), and the top 50 highly expressed genes (HEGs). C, Representative spatial gene-expression predictions for held-out invasive ductal carcinoma (IDC), pancreatic adenocarcinoma (PAAD), and colon adenocarcinoma (COAD) samples. H&E images, ground-truth spatial transcriptomic measurements, and predictions from ST-Net, HEST, BLEEP, HisToGene, Hist2ST, and GenVinci-2DST are shown. **D,** Construction of the adult aging mouse virtual transcriptomic atlas. GenVinci-2DST was applied to 63 H&E sections from young and old mice to reconstruct approximately 11.7 million predicted cells across 15 major organs; UMAPs show a random subsample of 100,000 reconstructed cells colored by age group, sample, and organ. **E,** Experimental validation of GenVinci-2DST predictions against STARmap measurements in representative old tongue and kidney sections. H&E images, STARmap measurements, and GenVinci-2DST predictions are shown for representative marker genes; measured and predicted expression were aggregated and smoothed within 100-μm windows (Methods). Right top, distribution of gene-level PCCs across the top 50 HVGs for old and young tongue and kidney samples. Right bottom, agreement between STARmap- and prediction-derived Leiden clustering (resolution 0.5; number of clusters not fixed), quantified by the adjusted Rand index (ARI). **F,** Senescence-associated transcriptional landscape of the reconstructed aging mouse atlas. Left, organ-level heatmap of SenNet-100, SenMayo, the two SenNet-100 modules, and the canonical senescence markers p21 and p16. Right, correlation between the SenNet-100 score and each marker within each organ.

We first evaluated GenVinci-2DST using held-out spatial transcriptomic datasets spanning five cancer types and two experimental platforms (Methods). Whereas existing methods, including ST-Net^13^, HEST^32^, BLEEP^22^, HisToGene^20^, and Hist2ST^21^, were retrained separately on each dataset using leave-one-sample-out cross-validation, GenVinci-2DST was trained once on paired profiles in ImageOmicsNet-300M and subsequently evaluated in a zero-shot setting without any dataset-specific retraining or fine-tuning. Despite this substantially more challenging evaluation setting, GenVinci-2DST achieved the highest average prediction accuracy across the top 50 highly variable genes (HVGs), the top 50 highly expressed genes (HEGs), and all genes, reaching average PCCs of 0.45, 0.42, and 0.33, respectively, versus 0.35, 0.33, and 0.22 for HEST, the strongest baseline on average (Figure 4B and Figure S5C). Spatial visualization showed that GenVinci-2DST recovered fine-scale gene expression patterns and delineated tumor and normal regions in held-out cancer samples (Figure 4C and Figure S5D), further supporting generalization across unseen donors. Model interpretability and uncertainty were further examined on a representative breast invasive ductal carcinoma (IDC) section (Methods). Leave-one-sub-patch-out occlusion analysis revealed a positive association between bin-level predictions and their contributions to spot-level expression for every gene examined (mean Pearson *r* = 0.45), and this agreement varied with tissue morphology, being highest in stromal regions for the fibroblast marker *POSTN* (Figure S6A). Test-time augmentation (TTA)-derived uncertainty tracked prediction error both across genes (median Spearman ρ = 0.18; 98.0% of genes showed positive correlations) and across spatial locations (Spearman ρ = 0.73), with the least confident patches corresponding to sparse, structurally uninformative tissue (Figure S6B).

Next, we asked whether GenVinci representations could be transferred at the multi-organ scale. We applied GenVinci-2DST to 63 H&E sections spanning 15 major organs from young and old mice, generating a multi-organ virtual transcriptomic atlas of approximately 11.7 million predicted cells (Figure 4D and Figure S6C). For validation, we additionally profiled sections by STARmap and then H&E-stained the same sections—a stringent test, with tongue absent and kidney comprising only 1% of the training corpus, and STARmap unseen during training. All predictions in the atlas and in the validations above were generated zero-shot, without fine-tuning on STARmap or on any mouse tissue. At 100 × 100 µm, the scale at which the model is supervised during pretraining, GenVinci-2DST recovered the spatial expression patterns of representative marker genes and HVGs in both organs, with individual genes reaching PCCs of up to 0.82 (Figure 4E and Figure S6D,E). Mean PCCs across the top 50 HVGs ranged from 0.11 to 0.16, a range consistent with a targeted 890-gene panel that excludes the highly expressed transcripts most readily predicted from histology^53^. More importantly, the reconstructions preserved higher-order spatial organization: independent clustering of predictions and measurements yielded ARIs of 0.57–0.72 and NMIs of 0.78–0.90, recovering comparable numbers of spatial domains in both modalities (Figure 4E and Figure S6F–H). Correspondence was lower at bin-level resolution (mean PCC 0.05–0.08), reflecting both the sparsity of the ground truth at this scale, where windows contain a median of only 220–380 transcripts, and the absence of direct bin-level supervision during pretraining, where the loss constrains only the spot-level mean of the 8 × 8 bin outputs. That this reflects a measurement-limited ceiling rather than a failure of transfer is indicated by models trained directly on STARmap: fine-tuning on the target organ reached a mean PCC of only 0.10 across four cross-sample configurations, and a model trained from scratch on the H&E embedding alone 0.08, compared with 0.05 for zero-shot prediction (Figure S6I). As an independent validation, predicted expression of representative neuronal, glial, and regionally expressed markers in young and old mouse brain recapitulated the major anatomical patterns of the Allen Brain Atlas in situ hybridization (ISH) references^54^ (Figure S6J).

We next examined whether this fidelity extended beyond individual genes to coordinated senescence programs. In a representative old mouse tongue sample, H&E-inferred SenNet-100 showed spatial distributions corresponding to those of the canonical senescence markers *Cdkn2a* (p16) and *Cdkn1a* (p21), with all three enriched in the lingual epithelium (Figure S7A). We then extended the senescence analysis across the adult mouse atlas (Figure 4F and Figure S7B). Because *Cdkn1a* and *Cdkn2a* were excluded from all signatures, they provide an internal consistency check that is not driven by gene overlap: within individual organs, the SenNet-100 score tracked p21 at cellular resolution (median Spearman ρ = 0.71; ρ > 0.5 in 13 of 15 organs), whereas p16 was far weaker (median ρ = 0.18), consistent with its sparse expression (Figure 4F, right, and Figure S7B). Senescence differed substantially across organs, and this ordering was reproduced by independent signatures (SenMayo ρ = 0.70, *P* = 3.7 × 10⁻^³^; p21 ρ = 0.91, *P* = 3.1 × 10⁻^⁶^; Figure 4F, left), with the two SenNet modules dissociating by tissue class—the stress/inflammatory module dominating in lymphoid organs and cytoskeletal/adhesion in stratified epithelia (Figure S7C). Brain was the sole exception on every internal-consistency metric, likely reflecting limited morphological information content rather than low senescence (Figure S7D). Age-associated differences were small by comparison and only moderately concordant between signatures (ρ = 0.56, *P* = 0.031; Figure S7E,F); as each organ was represented by one old and one young animal, these differences are reported descriptively. Nevertheless, these analyses show that virtual transcriptomics reconstructed from histology retains senescence-associated programs at cellular resolution and resolves their organ-specific composition.

### Serial histology extends virtual spatial transcriptomics to cellular-resolution 3D tissue scenes

We next asked whether GenVinci could extend histology-derived transcriptomic reconstruction from two-dimensional sections to 3D tissue architecture. We collected human abdominal skin samples from nine female donors spanning six ages (20, 23, 32, 40, 51, and 71 years), and generated 1,719 serial H&E sections at 10-µm thickness. For each donor, 100 anatomically matched consecutive sections were manually selected, of which 605 (45–81 per donor) passed manual quality control to exclude tissue artifacts, scanning artifacts, and blurred images. GenVinci-3DST was then applied for tissue segmentation, rigid and non-rigid registration^55^, spatial gene prediction, and downstream analysis (Methods) to generate 3D virtual spatial transcriptomic profiles, enabling reconstruction of 3D gene expression, cell-type organization, multicellular niches, and aging-associated molecular states from routine histology images (Figure 5A).

**Figure 5.**
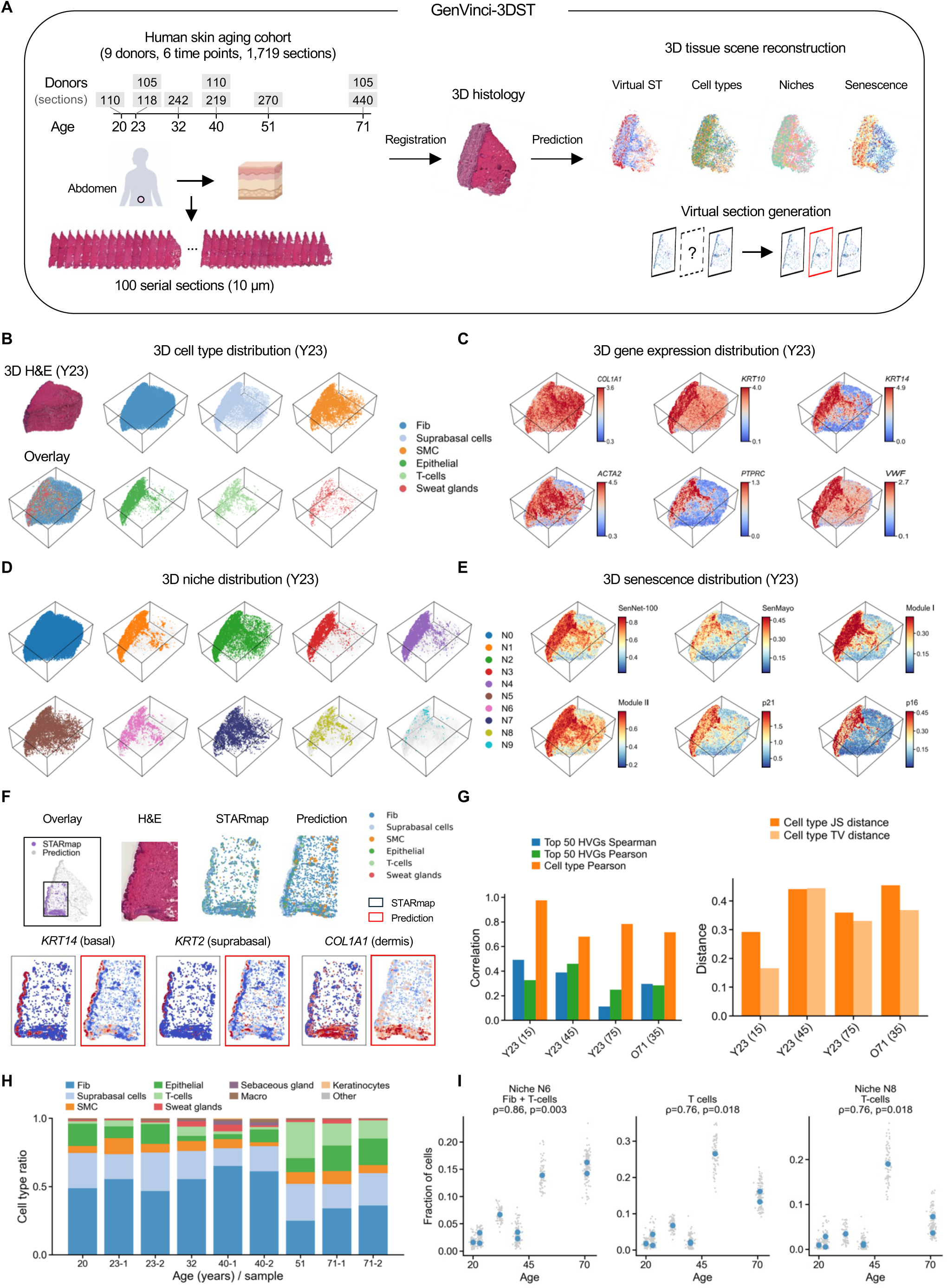
GenVinci-3DST reconstructs three-dimensional tissue scenes of human skin. **A,** Overview of the human female abdominal skin cohort (9 donors, 6 age time points) and the GenVinci-3DST workflow. Serial 10-μm sections (100 per donor) were registered into a three-dimensional histology volume; 605 sections (45–81 per donor) passed quality control. From the registered volume, virtual spatial transcriptomics, cell types, niches, and senescence distributions were predicted. Donors are labeled by age, with a numeric suffix distinguishing donors of the same age (23-1, 23-2, 40-1, 40-2, 71-1, 71-2); Y23 and O71 in B–G refer to donors 23-1 and 71-1. **B,** Left, reconstructed three-dimensional H&E volume of donor Y23. Right, three-dimensional distribution of each major cell type, shown in a shared view and bounding volume. **C,** Three-dimensional distribution of predicted expression for dermal (*COL1A1*), epidermal (*KRT10*, *KRT14*), smooth-muscle (*ACTA2*), immune (*PTPRC*), and endothelial (*VWF*) markers. Color scales are clipped at the 1st and 99th percentiles and are computed separately for each donor. **D,** Three-dimensional distribution of the ten cellular niches (N0–N9), defined by clustering local cell-type composition across all nine donors (Methods): N0, fibroblast; N1, suprabasal; N2, suprabasal–fibroblast; N3, epithelial; N4, fibroblast–epithelial; N5, fibroblast–smooth muscle; N6, fibroblast–T cell; N7, smooth muscle; N8, T cell; N9, macrophage. **E,** Three-dimensional senescence distribution: SenNet-100 and its stress–inflammatory (Module I) and cytoskeletal–adhesion (Module II) modules, SenMayo, and p21 (*CDKN1A*) and p16 (*CDKN2A*) expression. *CDKN1A* and *CDKN2A* were excluded from all gene sets and serve as independent axes. Color scales are clipped at the 2nd and 98th percentiles and are shared between donors. **F,** Validation against a STARmap section adjacent to Y23 section 15. Top left, spatial overlap of the profiled region; top right, H&E, STARmap, and predicted cell-type maps; bottom, paired STARmap and predicted expression for *KRT14* (basal), *KRT2* (suprabasal), and *COL1A1* (dermis). **G,** Quantitative agreement between prediction and STARmap for four matched sections (23-1 sections 15, 45, and 75; 71-1 section 35). Left, correlation of the top 50 HVGs and of cell-type composition; right, Jensen–Shannon and total-variation distances between cell-type compositions. **H,** Cell-type composition of each donor, ordered by age. **I,** Fraction of cells assigned to niche N6, to the T-cell class, and to niche N8 as a function of donor age (*n* = 9 donors). Blue, donor-level values; gray, individual sections. Spearman *ρ* and two-sided *p* are indicated.

In a representative young donor (age 23, Y23), the registration pipeline progressively improved alignment across adjacent H&E sections and produced a coherent 3D morphology of human skin (Figure 5B, left; Figure S8A,B). Based on the reconstructed volume, GenVinci-3DST recovered the major cellular compartments in three dimensions, including fibroblasts, suprabasal cells, smooth muscle cells, epithelial cells, T cells, and sweat gland cells (Figure 5B, right). Cell-type proportions remained largely continuous across serial sections, supporting the spatial consistency of the reconstructed atlas (Figure S8C, left). The atlas further preserved spatially organized gene expression programs, with canonical markers showing compartment-specific localization across the volume: dermal (*COL1A1*), epidermal and basal (*KRT10*, *KRT14*), smooth muscle (*ACTA2*), immune (*PTPRC*), and endothelial (*VWF*)^56^ (Figure 5C). Beyond individual genes and cell types, GenVinci-3DST resolved ten coherent multicellular niches defined jointly across all nine donors, including fibroblast–smooth muscle (N5), fibroblast–epithelial (N4), and fibroblast–T cell (N6) niches, together with a continuous epidermal–dermal interface niche (N2) (Figure 5D; Figure S8C, middle). The reconstructed atlas also enabled spatial characterization of senescence programs, which formed localized domains within the volume rather than a uniform elevation (Figure 5E; Figure S8C, right). Similar 3D morphology, cell-type organization, gene expression patterns, niche structure, and senescence landscape were observed in an old donor (age 71, O71), supporting reproducibility across age groups (Figure S8D–H).

We validated the predicted profiles against matched STARmap measurements from four adjacent sections, three from the Y23 donor and one from the O71 donor (Methods). Predicted slide-level profiles recapitulated STARmap measurements, with strong agreement in cell-type composition (mean PCC = 0.79, range 0.68–0.98; mean Jensen–Shannon distance = 0.39; mean total-variation distance = 0.33) and positive concordance in gene expression across the top 50 HVGs (mean Spearman ρ = 0.32; mean PCC = 0.33), while canonical markers reproduced the spatial localization of basal, suprabasal, and dermal compartments (Figure 5F,G; Figure S9A). Senescence scores were consistent with canonical markers within the reconstructed volumes: SenNet-100 showed substantially greater concordance with p21 (*CDKN1A*) than did the SenMayo signature in both donors (Spearman ρ = 0.79 and 0.84 vs. 0.63 and 0.72), with weaker but concordant associations with p16 (*CDKN2A*); because *CDKN1A* and *CDKN2A* were excluded from all gene sets, these comparisons are not driven by shared genes (Figure S9B). GenVinci further generated virtual sections at unsampled depths that preserved cell-type organization and spatial gene expression continuity. Virtual sections matched the adjacent STARmap measurements as closely as the flanking predicted sections did (mean Spearman ρ = 0.33 vs. 0.32 and 0.34) and recovered cell-type composition equally well (mean PCC = 0.82; mean Jensen–Shannon distance = 0.40), supporting the ability of the reconstructed atlas to infer intermediate tissue states along the 3D axis (Figure S9C,D).

We next asked what the third dimension resolves beyond individual sections. Multicellular niches that appeared as many separate patches within single sections were continuous in three dimensions. For example, the largest connected component of the epidermis-associated niche contained 86.9% (Y23) and 75.1% (O71) of its cells and spanned all sections, whereas the same cells were fragmented into 26–67 disconnected components within each section (Figure S10A). Moreover, redefining niches using two-dimensional neighborhoods reassigned 29% (Y23) and 43% (O71) of cells, with the mixed interface niches N4–N6 being least preserved (30–43% retained; Figure S10A). Thus, two-dimensional sections capture cell identity and overall composition but fragment continuous structures and under-resolve interfaces between compartments.

Finally, we leveraged the reconstructed 3D atlas across all donors to investigate age-associated cellular and molecular remodeling. Overall cell-type composition was broadly stable with chronological age, with fibroblasts and epidermal populations remaining the dominant compartments (Figure 5H). The clearest age-associated change was immune: the T-cell fraction rose from 1.3–6.8% in donors aged 40 years or younger to 13.3–26.5% in donors aged 51 years or older (Spearman ρ = 0.76, *P* = 0.018), accompanied by coordinated expansion of both T-cell-associated niches, N8 (T cells; ρ = 0.76, *P* = 0.018) and N6 (fibroblast–T cell; ρ = 0.86, *P* = 0.003), the latter showing the strongest association after Benjamini–Hochberg correction (FDR = 0.069; Figure 5I; Figure S10B). In contrast, whole-skin senescence scores showed no uniform age-dependent shift (|ρ| ≤ 0.28, all *P* > 0.4). An exploratory analysis within individual cell types instead revealed opposing, cell-type-specific trends: fibroblasts showed a relative increase in senescence-associated activity, tracked concordantly by SenNet-100, SenMayo, and p21 (*CDKN1A*) (ρ = 0.48, 0.58, and 0.63), consistent in direction with previous reports^57,58^, whereas T cells showed the strongest coordinated decline across all scores and both markers (ρ = −0.41 to −0.71; Figure S10C,D); these cell-type-specific trends require further validation.

These results demonstrate that GenVinci enables scalable reconstruction of 3D virtual spatial transcriptomes from serial H&E sections, recovers continuous tissue structure that individual sections cannot resolve, and provides a framework for characterizing cell-type-specific molecular remodeling during human tissue aging.

### Reconstructed tissue scenes reveal enrichment of genetic risk within disease-relevant spatial microenvironments

Having established that GenVinci captures biologically meaningful spatial representations, we next asked whether it could link tissue morphology to genetic susceptibility to disease. We therefore integrated GenVinci with gsMap^31^, which combines GWAS summary statistics with spatial transcriptomics to localize the cells through which genetic risk acts. Because gsMap was designed for sparse measured data, we introduced two modifications: GenVinci representations replace the PCA embeddings used as input to its graph-based variational step, and a magnitude-aware gene-specificity formulation accommodates dense expression profiles (Figure 6A). Together, these extend spatial genetic mapping to both imputed spatial transcriptomics (ST-only) and H&E-derived virtual transcriptomics (H&E-only).

**Figure 6.**
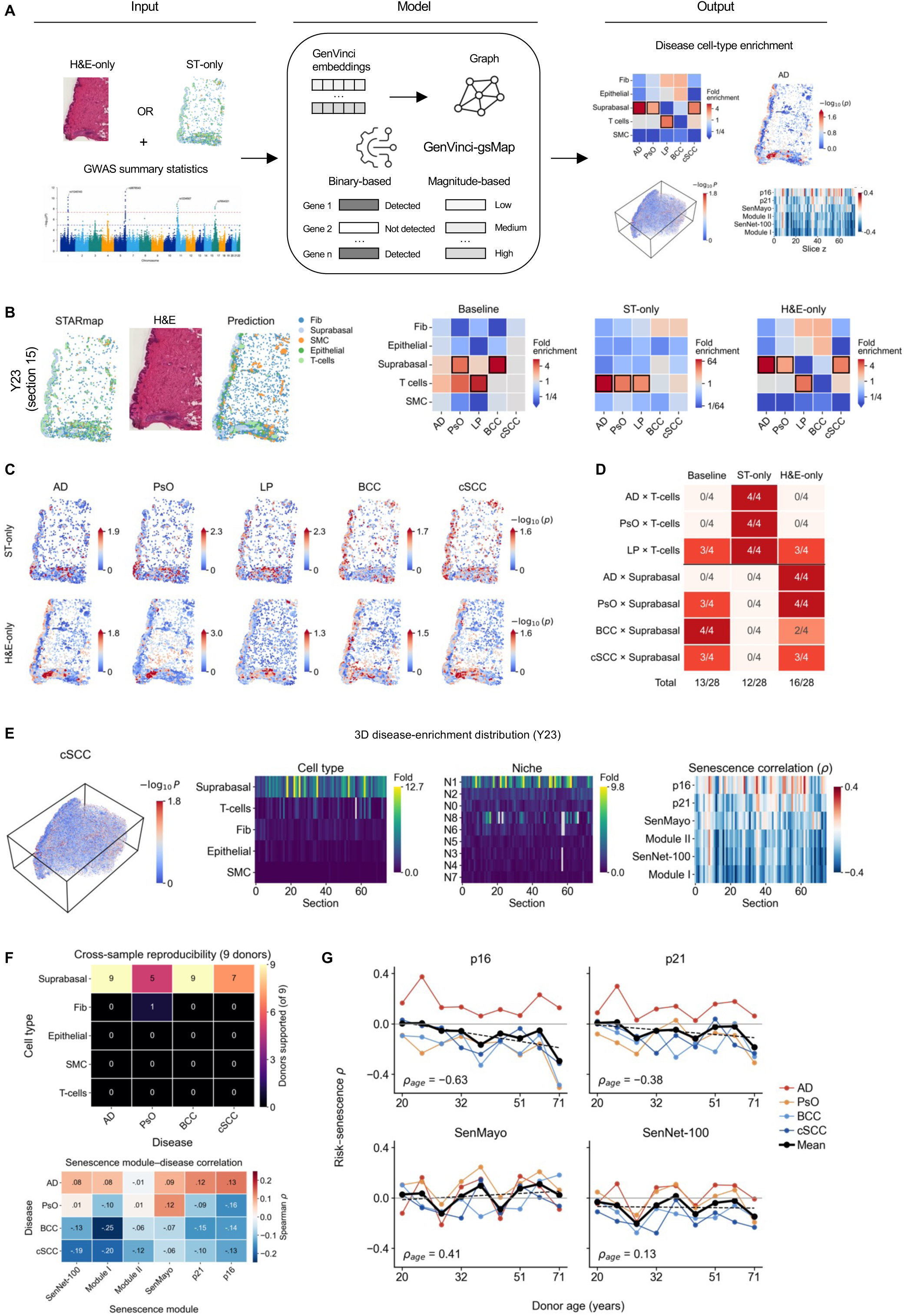
GenVinci-gsMap enables spatial mapping of disease-associated genetic susceptibility from measured and virtual transcriptomics. **A**, Overview of the GenVinci-gsMap framework. GWAS summary statistics are integrated with either measured spatial transcriptomics (ST-only) or H&E-derived virtual spatial transcriptomics (H&E-only). GenVinci representations replace the PCA input to the graph-based step of gsMap, and a magnitude-aware gene-specificity score enables analysis of imputed or predicted expression. **B,** Spatial genetic mapping in a representative section (Y23, section 15). Left, STARmap cell-type map, H&E, and predicted cell types. Right, disease–cell-type enrichment for five dermatological diseases with raw STARmap analyzed with the original gsMap (Baseline), imputed STARmap analyzed with GenVinci-gsMap (ST-only), and H&E-derived profiles analyzed with GenVinci-gsMap (H&E-only). Black boxes, literature-supported associations meeting all criteria (Methods). **C,** Per-cell association maps for the same diseases under the ST-only and H&E-only settings. **D,** Number of sections (of four) in which each literature-supported association was supported with each input. **E,** Three-dimensional disease-enrichment distribution for cSCC across 75 serial sections of Y23. Heatmaps show the enrichment of each cell type and niche, and the correlation between the per-cell association and each senescence measure among suprabasal cells, along the tissue-depth axis. **F,** Top, number of donors (of nine; 605 sections) in which each cell-type–disease association was supported. Bottom, correlation between the per-cell association and each senescence measure among suprabasal cells (median across donors). **G,** The same correlation as a function of donor age; thick line, mean of the four diseases. Color scales are independent per panel. Niches: N0, fibroblast; N1, suprabasal; N2, suprabasal–fibroblast; N3, epithelial; N4, fibroblast–epithelial; N5, fibroblast–smooth muscle; N6, fibroblast–T cell; N7, smooth muscle; N8, T cell. N9 (macrophage) was excluded (fewer than 30 cells in most sections). Disease abbreviations—AD, atopic dermatitis; PsO, psoriasis; LP, lichen planus; BCC, basal cell carcinoma; cSCC, cutaneous squamous cell carcinoma.

We analyzed five dermatological diseases—atopic dermatitis (AD), psoriasis (PsO), lichen planus (LP), basal cell carcinoma (BCC), and cutaneous squamous cell carcinoma (cSCC)—with mean corpuscular volume (MCV) and schizophrenia (SCZ) as negative-control traits (Table S3). Three inputs were compared on the four skin sections from two donors: raw STARmap (1,000 genes), imputed STARmap (approximately 16,000 genes), and virtual transcriptomics predicted from adjacent H&E sections (19,474 genes). For each cell type, we computed the fold enrichment of trait-associated cells relative to the rest of the section, and considered an association supported when it exceeded that of both control traits in the same cell type (Methods). We evaluated seven literature-supported associations forming two axes: an immune axis linking AD, PsO, and LP to T cells^59–63^, and an epidermal axis linking AD, PsO, BCC, and cSCC to suprabasal cells, the differentiated epidermal keratinocyte population^60,62,64–67^.

We first asked whether the measured data were sufficient to recover these associations. Raw STARmap did not resolve the immune axis: for AD, 31–47% of all cells in each section were classified as trait-associated irrespective of cell type, and neither AD nor PsO was associated with T cells in any section (Figure 6B,D; Figure S11B). This section-wide elevation was removed by imputation, after which only 1.5–2.2% of cells remained AD-associated and these were concentrated in T cells (52 of 86 T cells in the O71 section, compared with 1.7% of all cells). AD, PsO, and LP were consequently each associated with T cells in all four sections, with 22–150-, 11–43-, and 12–22-fold enrichment, respectively (Figure S11A). The limited resolution of raw STARmap therefore reflects the sparsity of the 1,000-gene panel rather than an absence of genetic signal in the tissue.

We then examined whether the same mapping could be achieved from histology alone. H&E-derived virtual transcriptomics recovered the epidermal axis in the absence of any measured expression: AD and PsO were associated with suprabasal cells in all four sections (3.9–14.0- and 1.8–4.7-fold enrichment), cSCC in three, and BCC in two (Figure 6B,D). These associations were disease-specific rather than a general property of the predicted profiles, as both negative-control traits were depleted in the same cells (MCV, 0.01–0.27-fold; SCZ, 0.02–0.48-fold; Figure S11A). Per-cell maps localized the trait-associated cells predominantly to the epidermal compartment (Figure 6C; Figure S11D). The H&E sections analyzed here are adjacent to the profiled STARmap sections; the two modalities therefore cannot be compared at the level of individual cells, and the agreement we assess below is at the level of cell types.

The associations recovered from H&E were concordant with the measured data. Those of PsO, BCC, and cSCC with suprabasal cells were likewise supported by raw STARmap (3, 4, and 3 of 4 sections), and that of LP with T cells was supported by all three inputs. Across the 28 association–section combinations, 16 were supported by H&E-derived input, compared with 13 by raw and 12 by imputed STARmap (Figure 6D); the two dense inputs thus recovered complementary axes, with imputation resolving the immune axis and H&E-derived profiles the epidermal axis. The choice of representation affected these results far less than the input data itself: substituting GenVinci representations for PCA left the number of supported associations unchanged, while increasing the enrichment of AD and PsO in T cells on imputed data by 19–94% in three of four sections (Figure S11C). Virtual transcriptomes derived from routine H&E thus recover literature-supported disease–cell-type associations, which motivated extending the analysis beyond the sections with matched measurements.

In reconstructed three-dimensional volumes, the epidermal axis persisted across consecutive sections (Figure 6E; one representative disease shown, remaining diseases in Figure S12A,B). For cSCC in the Y23 volume, suprabasal cells were the most enriched cell type in 71 of 75 sections and the association was supported in 74, forming a continuous band through the full depth of the tissue that is not resolvable from any single section. The same organization held at the level of spatial niches, where the suprabasal niche N1 was the most enriched niche in 59 of 75 sections. To test whether these patterns generalize, we applied the framework to all nine donors (605 sections). The four epidermal associations were reproducible across donors: AD and BCC were supported in the majority of sections in nine of nine donors, cSCC in seven, and PsO in five (565, 480, 441, and 325 of 605 sections, respectively), whereas no other cell type was supported for any disease in more than one donor (Figure 6F, top; Figure S12C,E,F). Neither the strength of these associations nor the fraction of suprabasal cells showed a significant trend with donor age (|ρ| ≤ 0.35), indicating that the cellular localization of genetic risk is largely stable between 20 and 71 years.

Because four diseases with distinct pathogenesis converged on the same cell type, we asked whether they map to the same cellular state within it. Among suprabasal cells in H&E-derived profiles, the per-cell association with AD correlated positively with *CDKN1A* (p21) and *CDKN2A* (p16) expression in all nine donors (median ρ = 0.12 and 0.13), whereas the associations with BCC and cSCC correlated negatively with p16, p21, SenNet-100, and its stress–inflammatory module (median ρ = −0.10 to −0.25; negative in at least seven of nine donors); PsO showed no consistent direction (Figure 6F, bottom). The two keratinocyte carcinomas and the inflammatory disease thus occupied opposite ends of the same axis, with carcinoma risk favoring keratinocytes expressing lower levels of the cell-cycle inhibitors p16 and p21. Consistent trends were observed for BCC and cSCC in the sections with STARmap measurements (Figure S12D), although the sparsity of the measured data limits this comparison, and these within-cell-type relationships warrant further validation. Of the six senescence measures, only the coupling with p16 showed an age-related trend shared by all four diseases: it tended to become more negative in older donors (ρ = −0.35 to −0.81; ρ = −0.63 for their mean), whereas the other measures showed no trend that was consistent across diseases (Figure 6G), suggesting that genetic risk is increasingly borne by p16-low keratinocytes with age.

Together, these results establish GenVinci-gsMap as a framework for spatial genetic mapping from measured and histology-derived transcriptomics. Genetic risk for four skin diseases converged on suprabasal keratinocytes, a localization that was continuous through tissue depth, reproduced in most donors, and stable across five decades of age. Within this shared cell type, risk was partitioned along lines that follow disease pathogenesis rather than tissue anatomy, separating an inflammatory disease from keratinocyte carcinomas by the expression of cell-cycle inhibitors. That these patterns were recovered from virtual transcriptomes indicates that routine H&E images can carry sufficient information to localize genetic susceptibility to the cell populations, and potentially the cell states, through which it is thought to act.

## DISCUSSION

In this study, we present GenVinci, a multimodal tissue foundation model that integrates transcriptomic representation learning with histology-based molecular reconstruction and 3D tissue scene modeling. Built on ImageOmicsNet-300M, a multimodal corpus of more than 300 million measured and H&E-predicted cellular and spatial profiles, GenVinci learns transferable representations of molecular state, cellular identity, and tissue context. These representations generalized across tissues, species, platforms, and modalities, suggesting that joint training on dissociated and spatial measurements can provide a common basis for interrogating tissue organization.

These representations transferred across tasks probing distinct aspects of tissue biology: GenVinci outperformed single-cell and spatial foundation-model baselines in cell-type annotation, spatial-domain identification, and batch integration, with the largest gains on a held-out platform and tissue. Cortical layers could be recovered without spatial coordinates, suggesting that transcriptional states themselves encode programs associated with tissue architecture. GenVinci embeddings also improved age prediction and senescence identification, tasks not targeted during pretraining.

At atlas scale, predicted senescence was organized primarily by cell identity, with vascular populations highest and naive lymphoid populations lowest across organs. Disease effects were bidirectional and often discordant between cell types within the same disease, so disease-level summaries can mask substantial remodeling. SenNet-100 partitioned, without functional input, into separable stress–inflammatory and cytoskeletal–adhesion modules. Although anchored on p16 and p21 detection, the classifier and SenNet-100 generalized beyond these markers in complementary ways: the classifier to experimentally induced senescence *in vitro*, and SenNet-100 to tissue *in vivo*, where senescent cells are rare and overlap with inflammatory states. These findings suggest that senescence is better represented as a context-dependent set of coordinated programs than by individual markers or a single fixed signature^29^, and that foundation-model representations can serve as a substrate for biological discovery rather than only as predictive features.

A second implication is that transcriptomic foundation models may provide molecular priors for translating morphology into molecular state. Trained once and applied zero-shot, GenVinci-2DST exceeded baselines retrained on each dataset. In tissues and a platform unseen during training, gene-level agreement was modest, but spatial domains agreed closely with measurements and predicted senescence programs tracked p21 within most organs. Virtual spatial transcriptomics therefore need not reproduce every transcript to recover biologically useful tissue states.

Across serial sections, GenVinci reconstructed cellular-resolution 3D tissue scenes of human skin that agreed with adjacent STARmap measurements, including at unsampled depths. In three dimensions, niches fragmented within single sections formed continuous structures, and interfaces between compartments, under-resolved by two-dimensional neighborhoods, were better resolved. Across donors, composition was broadly stable with age apart from an expansion of T cells and T-cell-associated niches, and senescence trends diverged between cell types, suggesting that tissue aging may manifest as remodeling of specific compartments and neighborhoods rather than as a global shift.

Integrating these tissue scenes with human genetics showed that dense expression, whether imputed or predicted from H&E, was required to resolve disease-relevant cell types that the sparse measured panel could not, whereas the choice of representation mattered far less than the input itself. The two dense inputs were complementary: imputation recovered an immune axis linking inflammatory diseases to T cells, and H&E-derived profiles an epidermal axis linking inflammatory diseases and keratinocyte carcinomas to suprabasal keratinocytes, likely reflecting the difficulty of resolving sparse immune cells from morphology and the limited within-compartment resolution of reference-based imputation. Across nine donors, the epidermal axis was continuous through tissue depth, reproduced in most donors, and stable with age, suggesting that the cell type through which inherited risk acts may be a stable property of the tissue. Within this shared cell type, however, risk for atopic dermatitis and for keratinocyte carcinomas tracked opposing levels of the cell-cycle inhibitors p16 and p21, and the coupling with p16 tended to strengthen with age. Carcinoma risk favoring keratinocytes with lower p16 and p21 is consistent with these loci acting in proliferation-competent cells, although differentiation state may contribute. These relationships were modest, derived from predicted expression, and supported only in trend by measured data; they are not causal but generate testable hypotheses linking inherited susceptibility, cellular state, and tissue organization.

### Limitations of the study

Several limitations remain. ImageOmicsNet-300M reflects biases in available data across tissues, species, diseases, and technologies. GenVinci does not encode spatial coordinates during pretraining; 3D reconstructions arise from registration and section-wise prediction. Without ablations under matched training data, the contributions of transcriptomic supervision, the spatial refinement module, and biological-context embeddings to histology prediction are not isolated, and gains over baselines may partly reflect pathology foundation-model features and the larger training corpus; we also did not compare against the most recent foundation models such as Novae^68^. Senescence mapping relies on weak supervision anchored mainly on p21 detection and an operational threshold; it may capture quiescence, differentiation, or stress programs, and requires orthogonal validation at the protein or SA-β-gal level. Histology-derived transcriptomes are predictions with modest accuracy at near-cellular scale, lower accuracy for sparse immune cells and lowly expressed genes, and sensitivity to slide-level staining, so weak transcriptional signals may be under-detected. Cell types in skin were assigned by reference-based label transfer, which depends on the reference. The aging analyses involved one animal per age group per organ and nine female donors from one sun-protected site, so age trends are exploratory. Genetic mapping relied on relative enrichment against two negative-control traits because absolute significance was not calibrated on dense inputs, was validated on four adjacent rather than identical sections from two donors, did not recover the immune axis from H&E, and identifies associations rather than mechanisms.

Overall, GenVinci connects molecular representation learning with virtual tissue reconstruction across cellular, spatial, and 3D scales. By transferring information from large transcriptomic atlases into histology, it enables tissue states to be inferred where direct multimodal measurements are unavailable, while retaining sufficient biological structure to study aging, senescence, and genetic susceptibility. Such virtual tissue models could complement experimental profiling by generating and prioritizing hypotheses at scales that are difficult to measure directly. Because H&E staining is inexpensive and widely available, these approaches could also extend molecular tissue analysis to clinical and resource-limited settings where spatial omics is impractical.

## Methods

### Construction of ImageOmicsNet-300M

#### Collection of public and consortium data

We constructed ImageOmicsNet-300M, a large-scale curated image–transcriptomics corpus comprising 300.2 million single-cell and spatial transcriptomic profiles and 4,623 H&E histology sections. The corpus contains 275.3 million experimentally measured single-cell and spatial transcriptomic profiles collected from over 20 public databases, the SenNet consortium (58.7 million cells), and more than 2,000 studies and publications, of which 6.3 million are paired with H&E histology across 2,378 sections, together with 24.9 million H&E-predicted virtual spatial profiles generated from 1,782 newly generated H&E sections, and 463 H&E images matched to spatial datasets from the SenNet consortium (Figure 1A and Table S1). ImageOmicsNet-300M spans two species (human and mouse), 82 tissue types, 55 transcriptomic technologies, and 1,420 annotated cell types.

For dissociated datasets, we collected approximately 89.6 million experimentally measured cells from public repositories, including CELLxGENE^35^ (85.8M), SenNet^34^ (3.7M), and additional datasets from GEO^69^ and other repositories (0.1M). These datasets cover diverse biological conditions, including development, aging, senescence, and disease.

For spatial transcriptomics data, we collected and generated approximately 210.6 million spatial profiles, including 185.7 million experimentally measured spatial profiles (including 66,956 newly generated STARmap cells) and 24.9 million H&E-predicted virtual spatial profiles. The experimentally measured spatial profiles were collected from multiple public resources and publications. Major sources included 10x Genomics (10.2M), SODB^33^ (33.1M), HEST-1k^32^ (2.1M), STimage-1K4M^36^ (4.2M), HuBMAP^70^ (1.4M), NanoString CosMx (3.0M), Vizgen MERFISH (12.3M), Allen Brain Map^71^ (9.0M), HCA^72^/HTAN^73^ (1.2M), SenNet^34^ (55.1M), and additional datasets manually curated from publications and consortium releases (53.9M). These resources span a broad range of spatial technologies, including Spatial Transcriptomics^3^, Visium, Visium HD^74^, Xenium^75^, MERFISH^4^, CosMx^76^, STARmap^5^, seqFISH^77^, and related targeted spatial assays.

Among these collected spatial datasets, 6.3 million profiles from HEST-1k and STimage-1K4M are paired with H&E histology images, corresponding to 2,378 paired histology sections. All HEST-1k and STimage-1K4M profiles were excluded in their entirety from foundation-model pretraining and reserved exclusively for histology-to-transcriptome training and evaluation, providing direct supervision for learning aligned representations between tissue morphology and spatial gene expression; together with the exclusion of the October 2026 SenNet release (see below), foundation-model pretraining therefore used 211.0 million experimentally measured profiles (Figure S1A).

SenNet data (58.7 million cells: 3.7M dissociated and 55.1M spatial profiles from 517 donors across 14 organs, including 463 spatial datasets with matched H&E images) were retrieved from the SenNet Data Portal^34^ (snapshot of October 16, 2026) and harmonized to AnnData following CELLxGENE schema conventions. This release supersedes the earlier SenNet data (0.8M) included in pretraining and was incorporated after model training; it was not used for GenVinci pretraining or any analysis in this study.

#### Newly generated data

To expand biological diversity and improve coverage of tissues relevant to aging and cellular senescence, we generated 1,782 H&E tissue sections for large-scale virtual molecular reconstruction. For the adult mouse aging atlas, we profiled 15 organs from six mice across young and old age groups, generating 63 H&E sections and 11.7 million predicted cells. For the human skin aging atlas, we collected 1,719 serial sections from abdominal skin samples obtained from nine female donors spanning six ages (20, 23, 32, 40, 51, and 71 years), yielding 13.2 million predicted cells and enabling 3D reconstruction of aging human skin.

For validation, we generated four additional same-section H&E–STARmap samples from mouse kidney and tongue, comprising one young and one old sample from each organ. For the human skin atlas, four sections within the serial-section series—three from a 23-year-old donor and one from a 71-year-old donor—were selected for STARmap profiling, while the surrounding serial sections were processed for H&E imaging. In addition, we generated an age-stratified human skin single-nucleus RNA-seq reference spanning five donor age ranges (20–29, 30–39, 40–49, 50–59, and 60–69 years). Each age group included multiple donors, and the combined dataset served as a pooled reference for cell-type label transfer to the human skin STARmap and virtual spatial transcriptomic datasets. We additionally generated an individual E14.5 mouse placenta STARmap sample for benchmarking representation-learning performance.

### Ethics statements

#### Animal studies

All animal experiments were approved by the Institutional Animal Care and Use Committee at Massachusetts General Hospital under protocol no. 2018N000182.

#### Human tissues

Human skin samples were derived from discarded healthy skin tissue obtained specifically for this study from abdominoplasty procedures performed at Massachusetts General Hospital (MGH). The use of discarded human tissue was reviewed and approved by the MGH Institutional Review Board (IRB; protocol 2020P001819). The IRB determined that the study did not meet the regulatory definition of human subjects research. Accordingly, informed consent for research participation and donor compensation were not required.

### Sample collection and tissue preservation

#### Adult mouse tissues

For adult mouse tissue samples, four 2-month- and two 26-month-old C57BL/6J mice (for young and old conditions, respectively) were euthanized in accordance with the approved IACUC protocol. The following tissues were immediately harvested: brain, heart, liver, spleen, pancreas, kidneys, dorsal skin, lungs, thymus, tongue, gastrointestinal tract (stomach, duodenum, jejunum, ileum, colon, and rectum), testes, arm muscle, leg muscle, and seminal vesicles. Each tissue was embedded in optimal cutting temperature compound (OCT) in a plastic mold, frozen on dry ice, and stored at −80 °C.

#### Human skin tissues

For the human skin serial H&E imaging and STARmap imaging, nine female donors spanning 20–71 years of age were included. Fresh-frozen tissue blocks were stored at −80 °C until sectioning and downstream processing.

#### Mouse placenta

One E14.5 placenta from a C57BL/6J × CAST/EiJ cross was used to generate the STARmap dataset. The placenta was embedded in OCT compound in a plastic mold, frozen on dry ice, and stored at −80 °C.

### Tissue sectioning and H&E imaging

#### Slide and plate preparation

For the mouse aging atlas, mouse kidney and tongue validation samples, and human skin samples, microscope adhesion slides were coated with poly-D-lysine for 2 h, washed three times with water for 5 min each, and air-dried in a fume hood for 2 h. For the individual E14.5 mouse placenta STARmap sample, glass-bottom six-well culture plates were activated by oxygen plasma and pretreated with γ-methacryloxypropyltrimethoxysilane (Bind-Silane), followed by the same poly-D-lysine coating procedure used for the microscope slides.

#### Cryosectioning

OCT-embedded mouse and human tissue blocks were cryosectioned at 10-µm thickness using a Leica CM3050 S cryostat. Sections from the mouse aging atlas and human skin aging atlas were mounted on the prepared microscope slides. Additional mouse kidney and tongue sections designated for same-section STARmap–H&E validation were prepared in the same manner. Human skin sections designated for STARmap validation were mounted separately and processed only by STARmap, whereas the individual E14.5 mouse placenta section was mounted in the prepared glass-bottom plate for STARmap analysis. All sections were stored at −80 °C until downstream processing.

#### H&E staining and imaging of atlas sections

H&E-only sections from the mouse aging atlas and human skin aging atlas were equilibrated to room temperature for 5 min and fixed in methanol for 5 min at −20 °C. Sections were stained using a Sakura Histo-Tek Slide Stainer, coverslip-mounted with Cytoseal XYL, and imaged using a Zeiss Axio Scan 7 slide scanner with a 20× objective lens at an effective pixel size of 0.172 µm × 0.172 µm. Images were processed using ZEN Blue software (Zeiss, Germany).

#### Same-section H&E imaging following STARmap

For the four mouse kidney and tongue validation sections, STARmap sequencing and imaging were completed before H&E staining. After the final STARmap imaging cycle, the imaging buffer was removed and the sections were washed three times with 1× PBS for 3 min each. The same tissue sections were then stained with H&E, coverslip-mounted with Cytoseal XYL, and imaged using the same whole-slide imaging workflow described above.

### STARmap in situ sequencing

#### Gene panel design

STARmap in situ sequencing (STARmap-ISS) was adapted from a previous protocol^5^ with modifications. For mouse kidney and tongue tissues, an 890-gene panel was designed, comprising 700 HVG genes and 190 aging- and senescence-related genes. For individual E14.5 mouse placenta tissue, we used a 903-gene panel that included both highly variable genes and placenta-specific markers. For human skin, a 1,000-gene panel was designed, comprising 825 canonical human cell-type marker genes and 175 aging- and senescence-related genes. All STARmap probes can be found in Table S4.

#### Fixation and permeabilization

Mouse kidney and tongue sections, the individual E14.5 mouse placental section, and human skin sections designated for STARmap-ISS were equilibrated to room temperature for 5 min and fixed with 4% paraformaldehyde (PFA) in 1× PBS for 15 min.

Sections were washed three times with PBSTR consisting of 0.1% Tween-20 and 0.1 U/µL SUPERase·In in PBS, followed by permeabilization with methanol at −20 °C for 30 min. Before probe hybridization, sections were rehydrated for 10 min in post-fixation buffer containing 10 mM Tris-HCl (pH 7.5), 1× PBSTR, and 0.1 mg/mL yeast tRNA.

#### Probe hybridization, ligation, and rolling-circle amplification

SNAIL probes were synthesized by Integrated DNA Technologies (IDT). SNAIL probes were resuspended in nuclease-free water and pooled according to tissue type. Before hybridization, probe mixtures were heated to 40 °C for 15 min and subsequently equilibrated to 37 °C. Sections were then incubated in hybridization buffer containing 2× SSC, 10% formamide, 0.1% Tween-20, 20 mM ribonucleoside vanadyl complex, 0.1 mg/mL yeast tRNA, 0.1 U/µL SUPERase·In, and SNAIL probes.

The final SNAIL probe concentration was 1 nM per probe for the individual E14.5 mouse placental tissue and 4 nM per probe for mouse aging atlas tongue, kidney, and human skin tissues.

Probe hybridization was performed at 40 °C with gentle shaking for 24 h for mouse placental tissue and 48 h for mouse kidney, tongue, and human skin tissues. Following hybridization, sections were washed twice with PBSTR for 20 min at 37 °C, followed by a 20-min wash with 4× SSC in PBSTR at 37 °C and a final rinse with PBSTR at room temperature.

Annealed SNAIL padlock probes were ligated in a T4 DNA ligation mixture containing a 1:10 dilution of T4 DNA ligase, 0.2 mg/mL BSA, and 0.5 U/µL SUPERase·In for 2 h at room temperature with gentle agitation. Sections were then washed twice with PBSTR for 5 min each.

Rolling-circle amplification was performed using a reaction mixture containing a 1:50 dilution of Phi29 DNA polymerase, 250 µM dNTPs, 20 µM 5-(3-aminoallyl)-dUTP, 0.2 mg/mL BSA, and 0.2 U/µL SUPERase·In. Samples were incubated for 2 h at 30 °C with gentle agitation and subsequently washed twice with PBSTR for 5 min each.

#### Hydrogel embedding and tissue clearing

Individual E14.5 mouse placental and human skin sections were treated with 25 mM acrylic acid NHS ester for 2 h at room temperature with agitation, rinsed once with PBST (0.1% Tween-20 in PBS), and incubated in monomer buffer containing 4% acrylamide, 0.2% bis-acrylamide, and 2× SSC for 15 min.

No. 2 micro cover glasses were treated with Gel-Slick Solution according to the manufacturer’s instructions. After removal of the monomer buffer, 30 µL of polymerization mixture containing 0.1% ammonium persulfate and 0.1% tetramethylethylenediamine in monomer buffer was added directly to each sample. Samples were immediately covered with Gel-Slick-treated coverglass and allowed to polymerize for 1 h at room temperature.

Tissue–gel hybrids were washed twice with PBST and cleared overnight at 37 °C in proteinase K digestion buffer containing 50 mM Tris-HCl (pH 7.5), 100 mM NaCl, 1% SDS, and 0.2 mg/mL proteinase K. The following day, samples were washed twice for 5 min with quenching solution containing 0.5 mM AEBSF in 1× PBST, followed by a final rinse in 1× PBST.

Hydrogel embedding and tissue clearing were omitted for kidney and tongue sections from the mouse aging atlas to preserve tissue morphology for subsequent same-section H&E staining and imaging.

#### Dephosphorylation

All sections were treated with dephosphorylation buffer for 1 h at 37 °C. Mouse kidney, tongue, and human skin tissues were incubated with a mixture containing a 1:100 dilution of Antarctic phosphatase, 0.2 mg/mL BSA, and a 1:10 dilution of Antarctic Phosphatase Buffer in water. The individual E14.5 mouse placental tissue was treated with a 1:100 dilution of shrimp alkaline phosphatase and 0.2 mg/mL BSA. Samples were subsequently rinsed with PBST.

#### Sequencing and imaging

Six sequencing cycles were performed following the STARmap-ISS protocol with tissue-specific modifications. Each cycle consisted of treatment with stripping buffer containing 60% formamide and 0.1% Triton X-100 for 5 min, followed by three 5-min washes with PBST and incubation with the appropriate sequencing mixture.

For human skin sections, the sequencing mixture contained 0.5 mg/mL BSA, 10 µM reading probe for all six cycles, 5 µM fluorescent decoding probe, and a 1:25 dilution of T4 DNA ligase. Samples were incubated overnight for 12 h at room temperature.

For mouse tongue and kidney samples, the sequencing mixture contained 0.2 mg/mL BSA, 10 µM reading probe for cycles 1–2, 20 µM reading probe for cycles 3–6, 5 µM fluorescent decoding probe, and a 1:25 dilution of T4 DNA ligase. The increased reading-probe concentration in later cycles was used to compensate for the absence of tissue clearing. Samples were incubated overnight for 12 h at room temperature.

For individual E14.5 mouse placental tissue, the sequencing mixture contained 0.2 mg/mL BSA, 10 µM reading probe for all six cycles, 5 µM fluorescent decoding probe, and a 1:25 dilution of T4 DNA ligase. Samples were incubated for 3 h at room temperature.

Following each sequencing cycle, samples were washed three times for 10 min in washing and imaging buffer containing 2× SSC and 10% formamide before imaging. DAPI staining was performed after the sixth imaging cycle for subsequent nuclear segmentation.

Images were acquired using a Leica TCS SP8 confocal microscope equipped with a 405-nm diode and a white-light laser. Mouse kidney and tongue sections were imaged using a 40× long-working-distance air objective (NA 0.6), individual E14.5 mouse placental sections using a 40× oil-immersion objective (NA 1.30), and human skin sections using a 20× long-working-distance air objective (NA 0.4).

#### STARmap image processing and transcript decoding

Images from successive STARmap sequencing cycles were registered across rounds, and fluorescent signals corresponding to rolling-circle amplification products were detected and decoded according to their six-round barcode sequences. Nuclei were segmented using the DAPI channel, and decoded transcripts were assigned to individual cells to generate cell-by-gene expression matrices. Low-quality cells and ambiguously decoded transcripts were removed before downstream analyses.

#### Imaging preprocessing and registration

Image tiles were first deconvolved with Huygens (v23.04). Images acquired across sequencing rounds were subsequently registered in three dimensions using a Fourier cross-correlation approach implemented with NumPy (v1.26.4) and SciPy (v1.13.1). A single sequencing round was selected as the reference, and each remaining round was aligned to this reference by estimating the translational displacement corresponding to the peak of the three-dimensional cross-correlation function.

#### Spot detection and barcode decoding

Following registration, image data were reformatted for analysis with Starfish (v0.2.2). Amplicon signals were identified using the BlobDetector algorithm with Gaussian kernels spanning σ = 0.5–10 voxels across 10 kernel sizes. To reduce low-intensity background detections, spots falling within the lowest 10–20% of local intensity maxima were filtered out. Barcode decoding was performed using the PerRoundMaxChannel method, in which the channel with the highest signal intensity was selected independently for each sequencing round to construct a six-round barcode. The resulting barcodes were matched against the reference codebook to assign gene identities. Spots with incomplete barcodes or sequences that did not correspond to a valid codebook entry were excluded.

#### Cell segmentation and transcript assignment

Cellular regions were approximated by spatially expanding the nuclear segmentation obtained from StarDist^78^. DAPI images were first segmented using the pretrained 2D_versatile_fluo model in StarDist (v0.9.1), and the centroids of the resulting nuclei were extracted. To generate an expanded foreground mask representing the approximate cellular area surrounding each nucleus, the nuclear segmentation mask was Gaussian-smoothed (σ = 10 pixels), binarized using Otsu thresholding, and further expanded by morphological dilation with a disk-shaped structuring element of 10-pixel radius. Nuclear centroids were then used as markers for seeded watershed segmentation within this expanded foreground mask, thereby partitioning the dilated regions into individual putative cell territories. Decoded transcripts were assigned to cells according to their spatial coordinates, and transcripts located outside the segmented cellular regions were excluded from downstream analyses.

### Human skin single-nucleus RNA sequencing

Human skin samples used for single-nucleus RNA sequencing were collected across five donor age ranges (20–29, 30–39, 40–49, 50–59, and 60–69 years). Each age group comprised tissue pooled from multiple donors within the corresponding age range.

#### Nuclei isolation

Single-nucleus isolation was conducted according to the 10x Genomics guide (CG000505) using the Chromium Nuclei Isolation Kit with RNase Inhibitor (PN-1000494). Briefly, human skin tissues were cut into 100-µm slices, and nuclei were isolated using the following steps. Lysis buffer was added to the tissue slices, and the tissues were dissociated with a plastic pestle until homogeneous. The mixture was then incubated on ice for 10 min. The dissociated nuclei suspension was passed through a nuclei isolation column by centrifugation. Nuclei were pelleted by centrifugation at 500 rcf at 4 °C for 3 min, followed by the removal of cell debris. The nuclei pellet was washed and resuspended in wash and resuspension buffer, and the suspended nuclei were kept on ice before loading into the 10x Chromium Controller.

#### Library preparation and sequencing

Approximately 10,000 nuclei per sample were loaded onto the 10x Chromium Controller, and single-nucleus libraries were generated using the Chromium Next GEM Single Cell Multiome ATAC + Gene Expression kit (10x Genomics). Both gene-expression and chromatin-accessibility libraries were generated and sequenced using a NextSeq 2000 sequencer with P4 XLEAP-SBS reagent kits (Illumina); however, only the gene-expression data were used in the present study. Chromatin-accessibility libraries were sequenced using 50 cycles for Read 1, 49 cycles for Read 2, 8 cycles for Index 1, and 24 cycles for Index 2. Gene-expression libraries were sequenced using 28 cycles for Read 1, 90 cycles for Read 2, and 10 cycles each for Index 1 and Index 2.

#### Single-nucleus RNA-sequencing data processing

Raw sequencing data from the human skin single-nucleus multiome libraries were processed using Cell Ranger ARC v2.0.2 (10x Genomics) with the refdata-cellranger-arc-GRCh38-2024-A reference genome and default parameters. Gene-expression and chromatin-accessibility libraries were jointly processed to obtain nucleus-level RNA expression matrices and ATAC-seq fragment files. Although both modalities were used for quality control, only the gene-expression modality was used for the downstream analyses presented in this study.

Chromatin-accessibility quality control was performed using ArchR (v1.0.3). Arrow files were generated with minTSS = 3 and minFrags = 1,000. Nuclei failing these ATAC-seq quality-control criteria were excluded from the corresponding gene-expression dataset so that downstream analyses were restricted to nuclei passing quality control in both modalities.

Gene expression matrices were analyzed using Seurat (v5.3.1). Ambient RNA contamination was corrected independently for each sample using SoupX (v1.6.2). Doublets were identified independently for each sample using scDblFinder (v1.24.10), and only nuclei classified as singlets were retained. RNA quality control retained nuclei with >200 detected genes, 300–50,000 total RNA counts, and <10% mitochondrial transcripts. Sample-specific upper cutoffs for detected gene numbers were manually determined from the QC distributions to exclude high-complexity outliers. Mitochondrial and ribosomal protein genes were removed before downstream dimensionality reduction and integration.

A total of 24,153 nuclei were retained after quality control. Gene-expression counts were normalized using Seurat’s NormalizeData function, and 3,000 highly variable genes were selected using FindVariableFeatures from merged objects. Expression values were scaled and subjected to principal component analysis. Batch integration across age-group/sample identities was performed using Harmony through Seurat’s HarmonyIntegration framework. The first 30 Harmony dimensions were used to construct the nearest-neighbor graph, perform Leiden-style graph-based clustering at a resolution of 0.3, and generate the UMAP representation. Cluster-specific marker genes were identified using Seurat’s FindAllMarkers, and clusters were manually annotated on the basis of canonical skin cell-type markers.

### GenVinci spatial and single-cell foundation model

#### Transcriptomic tokenization

Raw gene expression profiles are typically sparse, high-dimensional, and noisy, posing challenges for direct representation learning. To address this, we employ a transformer-based transcriptomic foundation model to encode single-cell and spatial transcriptomic profiles into compact latent representations. GenVinci is a BERT-like transformer^38,79^ pretrained on 211.0 million experimentally measured profiles from ImageOmicsNet-300M, enabling context-aware representation learning and robust transfer across diverse downstream biological tasks. Benefiting from the flexible self-attention mechanism, GenVinci can process transcriptomic data generated from diverse sequencing platforms without requiring explicit gene-order alignment across datasets. Large-scale pretraining further enables the model to capture biologically meaningful gene–gene dependencies and cellular programs that generalize across tissues, species, and modalities.

Because transformer models operate on sequential inputs, transcriptomic profiles are converted from tabular gene expression matrices into ordered token sequences using a binning-based value encoding strategy. We first retain only protein-coding genes present in a predefined vocabulary derived from the GenVinci pretraining corpus. The vocabulary contains 16,265 human–mouse homologous genes, 3,396 human-specific genes, and 5,361 mouse-specific genes, together with three special tokens (CLS, PAD, and MASK) and four context tokens encoding modality, assay type, species, and tissue type.

Gene expression values are then normalized by library size and log-transformed to improve comparability across cells. For gene *i* in cell *c*, the normalized expression value is defined as:

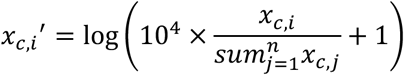

where *x_c_*_,*i*_ denotes the raw expression count, and *n* is the total number of genes. After normalization, genes with non-zero expression are identified for each cell. Their expression values are discretized using a binning-based value encoding strategy derived from scGPT^16^. Specifically, we use an even-width binning scheme with 50 expression bins, while an additional bin is reserved for zero-expression genes. Each gene is represented by two aligned tokens: a gene token, defined by the integer identifier of the gene in the predefined vocabulary, and an expression token, defined by its discretized expression bin. This encoding reduces sensitivity to small fluctuations in absolute expression values while preserving coarse expression magnitude information.

For each cell, a fixed-length sequence of 1,024 tokens is then constructed, with the first five positions reserved for CLS and context tokens. The remaining 1,019 positions are reserved for gene tokens. For cells with more than 1,019 non-zero genes, genes are sampled with probabilities proportional to their normalized expression levels. For cells with fewer than 1,019 non-zero genes, PAD tokens are appended to produce a fixed-length sequence. The gene-token embeddings for cell *c* are represented as:

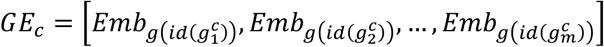

where *Emb*_g_ denotes a learnable embedding layer for gene tokens, *g^c^* denotes the *k*-th token in cell *c*, and *m* is the total sequence length. The aligned expression tokens described above are mapped to a separate learnable expression embedding layer, yielding expression embeddings (*EE_c_*). To further encode token categories, GenVinci also incorporates trainable segment embeddings (*SE_c_*), which distinguish different token types, including CLS, context and transcriptomic tokens. The final input embedding for each cell is defined as:

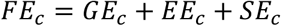

where *FE_c_* is an *m* × *d* matrix, and *d* denotes the embedding dimension. The resulting embeddings are then fed into stacked transformer blocks for contextual representation learning.

Multi-task masked-attention transformer pretraining

GenVinci uses a masked-attention transformer architecture to learn context-aware representations from paired gene and expression tokens. The model consists of 12 transformer encoder layers with a hidden dimension of 512, 8 attention heads, a feed-forward dimension of 2,048, GELU activation, layer normalization, and dropout rate of 0.10. For a given input embedding sequence, self-attention is computed as:

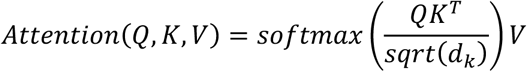

where *Q*, *K*, and *V* denote the query, key, and value projections derived from the input embeddings, and *d_k_* is the dimensionality of each attention head. Multi-head attention is then defined as:

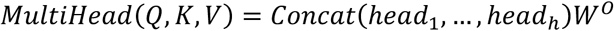

where 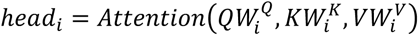, and 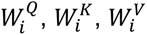 *W*^0^ are learnable projection matrices, and ℎ is the number of attention heads. The output of the attention module is followed by residual connection, layer normalization, and a position-wise feed-forward network:

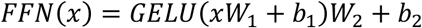

To support multi-task pretraining, GenVinci constructs two complementary masked views for each cell. In the gene-masked view, 15% of gene tokens are randomly masked while their corresponding expression tokens are retained. In the expression-masked view, 40% of expression tokens are randomly masked while gene identities are retained. Context tokens are kept unmasked in both views during training.

We pretrained GenVinci with three complementary objectives: masked gene prediction (MGP), masked expression prediction (MEP), and gene expression reconstruction (GER). MGP predicts the identities of masked gene tokens from the remaining gene, expression, and context information, encouraging the model to capture gene co-expression structure and gene-rank dependencies. MEP predicts masked discretized expression bins, encouraging the model to capture expression-state variation among cells with similar gene identities. GER reconstructs the normalized gene expression vector from the CLS representation, compensating for information loss introduced by token truncation and padding and the binning-based value encoding. The overall pretraining loss is defined as:

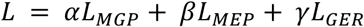

where *α*, *β*, and *γ* are task weights and are all set to 1 based on empirical tuning on the validation data. *L_MGP_* and *L_MEP_* are cross-entropy losses computed over masked gene and expression tokens, respectively, whereas *L_GER_* is a masked MSE loss computed between the reconstructed and normalized gene expression values over valid genes and cells. These three objectives jointly encourage GenVinci to learn robust cell-level representations that integrate gene identity, expression magnitude, and biological context.

### GenVinci senescence axis and module analysis

To investigate whether GenVinci embeddings capture biologically meaningful latent programs beyond canonical cell-type representations, we developed a weakly supervised framework to infer a global cellular senescence axis from pretrained embeddings. We analyzed the CELLxGENE-derived subset of ImageOmicsNet-300M, comprising 85.8 million cells with curated cell-type, condition, donor, and dataset information, spanning 930 cell types across 70 organs and 141 disease conditions in both human and mouse. Cell-type annotations were harmonized from source datasets and therefore retain differences in annotation granularity across studies.

Because detection of senescence markers depends strongly on factors unrelated to senescence (e.g., sequencing depth), seed labeling was performed within matched units rather than across the whole corpus. We therefore defined a stratum as species × cell type (1,240 strata), a donor unit as dataset × donor (16,889 combinations), and a depth decile as the within-stratum decile of the number of detected genes. Their intersection, termed a matching unit, is the unit within which every positive cell is matched to a control (1,109,071 matching units corpus-wide, of which 564,746 contain at least one matchable pair). Cells within a matching unit share species, cell type, dataset, donor, and comparable sequencing depth.

Human cells were labeled using *CDKN1A* (p21) and *CDKN2A* (p16), and mouse cells using their orthologs, *Cdkn1a* and *Cdkn2a*. Cells were labeled positive if either marker was detected and negative if neither was detected, without applying a continuous marker-score threshold. Positive cells were assigned to three evidence tiers: tier 1, both markers detected (0.93% of all cells); tier 2, p16 only (2.01%); and tier 3, p21 only (12.50%). This hierarchy prioritized double-positive cells, followed by p16-only and then p21-only cells, because p21 can also be expressed during transient stress, quiescence, and differentiation^80^. Within each stratum, positive cells were ranked first by tier, from tier 1 to tier 3, and then within each tier by the percentile of marker expression intensity calculated within their respective matching units. Following this ranking, each selected positive cell was paired with one negative cell from the same matching unit. Strata with fewer than 50 matchable pairs were excluded, and each retained stratum contributed at most 200 pairs to limit the influence of large strata. This procedure retained 1,022 of 1,240 strata and yielded 190,200 pairs (380,400 cells). Because double-positive cells were prioritized during selection, they constituted 34.6% of training positives despite accounting for only 0.93% of all cells in the original dataset. This strategy prioritized label confidence and matched comparisons over training set size.

Using these seed labels, we learned a global senescence axis in the pretrained GenVinci embedding space. For each seed cell, we extracted a 512-dimensional CLS embedding after removing the *CDKN1A* and *CDKN2A* tokens, or their mouse orthologs, from the input sequence. This masking prevented the classifier from directly accessing the genes used to define the labels. The embeddings were standardized and used to train a neural network with 512 input units, 256 hidden units, and one output unit to distinguish seed-positive from seed-negative cells. We then applied the same masking, standardization, and trained classifier to all cells in the atlas, assigning each cell a scalar probability score representing its relative position along the senescence axis. For downstream analyses, the top 5% of cells by score across the entire corpus (4,290,496 cells) were labeled as predicted senescent. This cutoff was used consistently as an operational definition and does not represent an estimate of the true prevalence of senescence.

To characterize senescence across biological contexts, scores were summarized across species, organ, cell type, and disease condition by computing both the fraction of predicted senescent cells and the mean score within each group, weighting groups by donor rather than by cell. For cell-type rankings (Figure 3B), cell types were included if, among healthy samples, they had ≥1,000 cells in each contributing species–tissue group and, in aggregate, ≥5,000 cells, ≥5 species–tissue groups and ≥5 donors (86 of 722 cell types).

To identify genes associated with the classifier, we compared cells with high and low model scores within matching units. Cells within each matching unit were further divided into tertiles of an immediate-early gene (IEG) score, calculated from the 50 genes most strongly associated with model scores in a preliminary analysis, the majority of which were IEGs. From matching units with sufficient cells, we sampled 6,000 spanning 872 strata; each matching unit × IEG tertile combination defined a pairing unit, yielding 18,000 pairing units matched for species, cell type, dataset, donor, and sequencing depth, with additional stratification to reduce differences in acute stress responses. Within each unit, we compared mean gene expression between cells in the top and bottom 15% of model scores. Expression differences were averaged across units, and significance was assessed using a sign test on the proportion of units with a positive difference, with Benjamini–Hochberg correction. Genes with FDR < 0.05 and a positive mean difference were retained. We then excluded IEGs, housekeeping genes, the seed markers *CDKN1A* and *CDKN2A*, S/G2M cell-cycle genes, and histone genes. The 100 remaining genes with the largest mean expression differences were selected as the SenNet-100 signature (Table S2).

To identify substructure within SenNet-100, we calculated pairwise Pearson correlations between genes using their expression differences across the 18,000 pairing units. The resulting 100 × 100 correlation matrix was subjected to hierarchical clustering. Cutting the dendrogram into four clusters yielded two principal modules containing 46 and 43 genes and two minor clusters containing 8 and 3 genes. Over-representation analysis was performed using Enrichr^81^ with the GO Biological Process 2023, GO Cellular Component 2023, Reactome 2022, and MSigDB Hallmark 2020 libraries, with the 23,869 genes detected in the corpus as the background.

For external evaluation, gene set scores for SenNet-100 and SenMayo were computed using *scanpy.tl.score_genes*, whereas marker-based readouts (p16, p21, and their union) were derived from log-normalized expression values. Gene symbols were matched case-insensitively to accommodate differences between human and mouse nomenclature; signature genes without a matching symbol in the target species (e.g., *IL32* in mouse) were omitted from scoring in that species. The framework is not restricted to the p16/p21-based seed labels used in this study and can accommodate alternative marker sets or phenotype-derived scores for seed definition.

### GenVinci-2DST framework for histology-to-transcriptome prediction

To enable robust prediction of spatial gene expression from histology images across heterogeneous experimental conditions, we developed GenVinci-2DST, a teacher-guided framework that leverages embeddings from both GenVinci and a pathology foundation model. In this knowledge-distillation setup, the GenVinci transcriptomic encoder acts as the teacher, and its embeddings supervise the image encoder. GenVinci-2DST was trained on a multimodal subset of ImageOmicsNet-300M containing approximately 6.3 million paired H&E image and spatial transcriptomic profiles across diverse assays, species, and tissues. All transcriptomic data were represented at the spot level, including native Visium and ST spots, and pseudo-spots generated by HEST from higher-resolution platforms such as Xenium and Visium HD^32^, to provide a consistent spatial unit for pairing with H&E images. Consequently, transcriptomic supervision was available exclusively at the spot level. GenVinci-2DST also includes a Spatial Refinement Module (SRM) to refine spot-level signals into latent sub-patch-level representations, enabling cellular-resolution gene expression prediction.

For each spot, an H&E image patch (224 × 224 pixels; 112 × 112 μm) centered on the spot was encoded using H-optimus-1^52^, a pathology foundation model trained on heterogeneous histological datasets spanning multiple tissues, studies, and imaging conditions. In addition to the global CLS embedding, we extracted 256 local visual tokens arranged in a 16 × 16 grid to retain fine-grained spatial information. The SRM aggregated these local visual tokens into 64 latent sub-patch-level tokens arranged in an 8 × 8 grid using average pooling. Each token was then projected into a 512-dimensional latent space using a multilayer neural network. Context information describing assay type, species, tissue, and biological condition was incorporated to account for dataset heterogeneity. Each context field was embedded using a shared special-token embedding table and then passed through a field-specific projection, with a separate learnable scaling parameter for each context type. During training, only the biological condition token was randomly replaced with an unknown-condition token with a probability of 0.3 to reduce overfitting to dataset-specific condition labels.

Global image context was incorporated by projecting the pathology model’s CLS embedding into the same 512-dimensional space and adding it to each sub-patch-level representation. The resulting representations were reshaped into an 8 × 8 grid and refined using a lightweight two-layer convolutional block to capture local spatial dependencies among neighboring sub-patches and promote spatially coherent latent representations. Two prediction heads were then applied to the refined sub-patch-level representations. The first was an expression head that predicted non-negative expression values for a unified panel of 25,030 genes using a Softplus output activation. The second was a teacher-alignment head that mapped each sub-patch-level representation into the 512-dimensional GenVinci embedding space. Spot-level embeddings and expression predictions were obtained by averaging the corresponding outputs across all 64 sub-patches and all 64 bins, respectively. The measured gene expression profile for each spot was encoded using GenVinci to obtain a transcriptomic teacher embedding. GenVinci-2DST was trained to align the spot-level image-derived embedding *z_ima_*_g*e*_ with the corresponding teacher embedding *z*_g*ene*_ using a cosine similarity loss^30^:

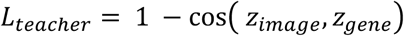

Gene expression supervision was applied using dataset-specific gene masks because different spatial transcriptomic datasets measure different gene panels. The overall training objective combined expression prediction loss and teacher distillation loss.

For inference, cell segmentation was first implemented on the whole-slide H&E image using the StarDist (*2D_versatile_he*) model^82^. Cell centroids were extracted from labeled segmentation masks, and a 224 × 224 H&E patch centered on each cell was cropped. The patch was passed through the pathology model encoder and GenVinci-2DST to generate an 8 × 8 grid of bin-level expression predictions. Although spot-level training used the average of all 64 bins, cell-centered inference used only the central 2 × 2 bins to cover the scale of the individual cell. The same central 2 × 2 averaging was applied to the predicted GenVinci embedding to obtain the prediction embeddings. This strategy reduces contamination from surrounding tissue and provides a localized near-cellular prediction while remaining consistent with spot-level supervision during training.

To assess whether predicted bin-level expression maps captured spatial contributions to the final spot-level predictions, we also performed a leave-one-sub-patch-out occlusion analysis. For each spot, each of the 64 image sub-patches was independently masked by setting its image embedding to zero while keeping all remaining inputs unchanged. The model was re-evaluated after each occlusion, and the contribution of each sub-patch was quantified as the decrease in the corresponding spot-level gene prediction after occlusion. The resulting occlusion importance map was compared with the predicted bin-level expression map using the Pearson correlation across the 64 sub-patches for every spot and gene. Gene-level statistics were summarized by the median correlation across all spots.

Predictive uncertainty was estimated using test-time augmentation (TTA). For each spot, nine translated views (a 3 × 3 grid of 0/±16-pixel shifts, including the unshifted image) were evaluated independently by the model. The mean prediction across augmented views was used as the final prediction, and the standard deviation across augmented predictions was used as the uncertainty estimate for each gene. Gene-wise uncertainty was validated by computing the Spearman correlation between prediction uncertainty and absolute prediction residual across all spots. To summarize uncertainty at the tissue level, uncertainty values for each gene were converted to within-gene percentiles before averaging across genes, yielding a spatial uncertainty map for each tissue section.

### GenVinci-3DST framework

#### Image registration and gene expression prediction

To reconstruct spatial transcriptomic landscapes across serial tissue sections, we developed GenVinci-3DST, an integrated framework that combines serial-section registration, cell segmentation, GenVinci-2DST, and virtual section generation. The framework takes serial H&E-stained tissue sections as input and generates virtual spatial transcriptomics in a shared registered coordinate system.

Serial H&E sections were first registered using the VALIS tool^55^. For each sample, section images were sorted according to slide index and registered in their physical order. VALIS was used to perform rigid and non-rigid registration across serial sections, and the warped images were saved using the overlapping tissue region as the final crop. This produced an aligned serial-section image stack for downstream 3D reconstruction.

Cell nuclei were segmented on each registered section using the same StarDist (*2D_versatile_he*) model. For each detected nucleus, the cell identifier and bounding box were saved, and the bounding-box center was used as the cell coordinate for patch extraction. Each patch was then sent to the trained GenVinci-2DST model for gene expression prediction. The same central 2 × 2 averaging was applied to both predicted gene expression and embeddings. The final GenVinci-3DST output is a registered serial-section collection of cellular-level predicted transcriptomics and embeddings, enabling 3D spatial analysis of tissue architecture and molecular organization.

#### Cell type and niche analysis

To assign cell types to predicted cells, we used the pooled snRNA-seq dataset from the matched reference age group. The reference dataset was scaled using precomputed gene-level means and standard deviations, followed by PCA and construction of a *k*-nearest-neighbor graph. Predicted cells were projected into the reference PCA space, and cell-type labels were transferred from the reference using a PCA-based kNN transfer strategy implemented with *scanpy.tl.ingest*. This produced a predicted cell-type annotation for each inferred cell.

For niche analysis, predicted cells were represented by their three-dimensional spatial coordinates and cell-type labels. We first harmonized the transferred labels into ten classes shared by all donors, including fibroblasts, suprabasal cells, T cells, epithelial cells, smooth muscle cells, sweat glands, sebaceous gland cells, macrophages, and keratinocytes, with the remaining rare labels grouped as other. Suprabasal cells and keratinocytes were retained as separate classes, as provided by the transferred labels. Spatial coordinates were expressed in micrometers, with the in-plane position taken from the registered section and the depth position from the section index and the 10-µm section thickness. For each cell, we identified its ten spatial nearest neighbors within the same donor, including the cell itself, and computed a local niche composition vector defined as the fraction of neighboring cells belonging to each retained cell class.

Local niche composition vectors were clustered using *k*-means clustering to identify recurrent spatial microenvironments. In contrast to clustering each donor separately, we clustered all donors jointly so that a single set of niche definitions applies to every donor and niche proportions can be compared across age. Each niche cluster was annotated according to the cell classes contributing at least one fifth of its centroid, listed in decreasing order of abundance, resulting in interpretable niche labels such as fibroblast–epithelial or suprabasal–fibroblast niches. We additionally quantified the enrichment of each cell class in each niche relative to its overall abundance across the cohort. The resulting niche identities were visualized in three-dimensional coordinates and summarized as per-section and per-donor compositions.

To assess what the third dimension contributes, we recomputed niche composition vectors using two-dimensional neighborhoods, in which the nearest neighbors of each cell were restricted to the same section, and assigned these vectors to the same niche centroids. Agreement between the two-dimensional and three-dimensional assignments was summarized as a row-normalized confusion matrix. We further assessed spatial continuity by connecting cells of the same niche within a fixed radius and extracting connected components, first across the whole reconstructed volume and then, for the same cells, within each individual section.

#### Virtual section generation

Since some tissue sections may be unavailable because they were reserved for external validation, excluded due to imaging artifacts, or failed quality control, we developed a spatial interpolation strategy based on GenVinci-3DST-predicted cell-level transcriptomics to generate virtual intermediate sections between adjacent registered tissue sections. For a pair of neighboring predicted sections, the cell coordinates from the left and right sections were first converted to the same raw image coordinate system. For each cell in the left section, neighboring cells in the right section were identified using a *k*-nearest-neighbor search. A spatially weighted estimate of the corresponding location in the right section was computed using a Gaussian kernel, and the virtual intermediate coordinate was defined as the midpoint between the original left-section coordinate and its estimated counterpart in the right section. In this study, 10 nearest neighbors were used for coordinate matching, with distance-based filtering to remove poorly matched cells.

Gene expression for each virtual cell was then generated by spatial kernel interpolation from the two adjacent predicted sections. For each virtual coordinate, nearby cells were retrieved independently from the left and right predicted sections. Their predicted expression profiles were aggregated using Gaussian spatial weights, and the left- and right-section estimates were combined with equal weighting to represent the middle section. For validation against measured intermediate STARmap sections, DAPI-based segmentation coordinates from the ground-truth section were transformed into the H&E coordinate system using the corresponding projective transformation matrix. The virtual section and measured STARmap section were then compared within their shared spatial coordinate range.

### GenVinci-gsMap framework

To perform spatial genetic trait mapping on GenVinci outputs, we integrated GenVinci with gsMap^31^, a framework that links genome-wide association study (GWAS) heritability signals to spatially resolved cellular microenvironments. The resulting framework, termed GenVinci-gsMap, extends the original gsMap to support transcriptomic embeddings and dense predicted transcriptomic profiles, enabling trait mapping from measured spatial transcriptomics, imputed transcriptomics, and H&E-derived virtual transcriptomics. Briefly, gsMap first learns a latent representation of each cell with a graph neural network variational autoencoder (GNN-VAE), defines a local microenvironment for each cell or spatial spot, computes gene-wise marker scores that capture local transcriptional enrichment, and then uses stratified linkage disequilibrium score regression (LDSC) to quantify whether genes enriched in specific spatial regions explain GWAS heritability^83^.

We introduced two modifications to the original gsMap framework to enable its application to both sparse spatial transcriptomic data and dense imputed or H&E-predicted virtual transcriptomics. First, we replaced the input features of the GNN-VAE, which in gsMap are principal components of the expression matrix, with GenVinci embeddings; the spatial graph, architecture, and training were unchanged. gsMap defines each spot’s microenvironment in two stages, first restricting candidates to spatially neighboring spots and then ranking these candidates by cosine similarity in the latent space; supplying GenVinci embeddings as input therefore preserves tissue context while grounding the latent space in features trained across tissues and modalities rather than derived from a single section.

Second, we introduced a continuous, magnitude-aware marker scoring mode to support dense transcriptomic matrices. The original gsMap was designed for sparse measured spatial transcriptomics and gates each gene by the ratio of local non-zero fraction to global non-zero fraction. However, this binary formulation becomes ineffective for imputed or predicted transcriptomics, in which the majority of genes are non-zero, the local and global fractions both approach one, and the resulting gate collapses every marker score to zero. To address this limitation, we computed the gate from the ratio of the mean expression within the local microenvironment to the mean expression across the entire tissue, which preserves the local-versus-global semantics of the original gate while remaining informative on dense data. The magnitude-aware mode was used for all inputs; on raw STARmap it gave near-identical results to the binary mode.

This generalized framework supports two practical scenarios. In the ST-only setting, imputed spatial transcriptomic matrices are combined with GenVinci embeddings to perform embedding-guided gsMap analysis. In the H&E-only setting, H&E images are first transformed into virtual transcriptomics and embeddings using GenVinci-2DST, after which the same embedding-guided, magnitude-aware gsMap framework is applied for disease-associated trait mapping.

Genome-wide expression for STARmap cells was imputed with Tangram^84^ by mapping the snRNA-seq reference dataset onto each section. The mapping was trained on the genes shared between the 1,000-gene STARmap panel and the reference dataset, and the learned mapping was used to project the expression of all reference genes onto the STARmap cells, yielding 15,907–16,162 genes per section. Imputed matrices were used without further filtering as the imputed STARmap input; raw STARmap counts (1,000 genes; 1.9–2.7% non-zero entries) were used as the raw STARmap input.

### Datasets for downstream tasks

We manually checked and made sure that all evaluation datasets were held out from pretraining. For the cell type annotation task, we benchmarked performance on one published MERFISH mouse brain dataset^40^ (40,153 cells × 1,085 genes across 10 cell types) and our newly generated mouse placenta STARmap dataset (10,833 cells × 787 genes, the subset of the 903-gene panel present in the GenVinci tokenizer vocabulary, across 7 cell types).

For the spatial domain identification task, we evaluated two samples from a Visium human brain dataset^41^ (sample 151509 with 4,789 spots × 33,538 genes and sample 151674 with 3,673 spots × 33,538 genes), sectioned across six cortical layers and white matter, which provide prominent spatial patterns and gold-standard annotations.

For the batch integration task, we used a MERFISH human developmental thalamus dataset^42^ (49,844 cells × 139 genes across 3 batches and 3 cell types) to evaluate cross-batch robustness.

For age prediction, we retrieved two MERFISH mouse brain datasets^23^, comprising coronal and sagittal sections. The coronal dataset includes mice spanning ages 3.4 to 34.5 months across 20 time points (3.4, 3.8, 4.3, 5.4, 6.6, 9.8, 12.9, 15.5, 15.8, 18.8, 19.8, 21.4, 23.5, 24.6, 26.7, 28.5, 30.9, 32.6, 33.2, and 34.5 months), whereas the sagittal dataset covers 6 time points (3.8, 6.6, 8.6, 19.8, 23.5, and 26.7 months). To improve computational efficiency, we randomly subsampled 10% of the cells and retained only the six most abundant cell types, each containing more than 10,000 cells. This preprocessing resulted in datasets of 126,851 × 300 cells-by-genes for the coronal sections and 75,125 × 300 for the sagittal sections.

For senescence identification, we retrieved two spatial transcriptomics datasets with established annotations from previous studies. Specifically, we used two spatial transcriptomic datasets, including a Visium dataset of mouse muscle tissue collected at 2 and 5 days following notexin-induced injury^43^ (1,792 spots × 33,217 genes) and a Stereo-seq dataset of human brain tissue comprising paired samples from patients with Alzheimer’s disease and age-matched healthy controls^44^ (13,062 cells × 49,971 genes). Following previous studies^27^, we focused on the two severe Alzheimer’s disease samples and their matched controls. For all datasets, we adopted the original study definitions to annotate senescent and non-senescent cells.

For senescence classifier and SenNet-100 module validation, we used four single-cell datasets with experimentally defined senescence labels, comprising two human in vitro systems and two mouse in vivo systems. The first in vitro dataset (13,575 cells × 32,738 genes) profiles primary human cells by Drop-seq and contrasts senescent with non-senescent cells^48^. The second (45,187 cells × 36,601 genes) profiles WI-38 human fibroblasts under multiple senescence-inducing conditions^49^, which we evaluated as three independent comparisons against the shared proliferating control (replicative senescence, irradiation, and etoposide arms). For the in vivo datasets, we used p16-FDR/mCherry-positive cells in *Kras*-driven murine lung tumors^51^ (8,712 cells × 20,404 genes) and SnC-enriched microglia in the murine brain^50^ (9,406 cells × 19,318 genes). For all datasets, we adopted the original study definitions to annotate senescent and non-senescent cells.

For spatial gene expression prediction, we evaluated the framework on held-out spatial transcriptomics datasets from HEST spanning five cancer types^32^: invasive ductal carcinoma (IDC; 4 samples and 35,715 spots), pancreatic adenocarcinoma (PAAD; 3 samples and 7,274 spots), colonic adenocarcinoma (COAD; 4 samples and 15,655 spots), rectal adenocarcinoma (READ; 4 samples and 8,407 spots), and axillary lymph nodes from patients with IDC (LYMPH; 4 samples and 18,582 spots). The IDC, PAAD, and COAD datasets were generated using the Xenium platform, whereas the READ and LYMPH datasets were profiled using 10x Visium.

For the GenVinci-2DST evaluation, the generated adult aging mouse atlas comprised 6 samples yielding 11.7 million predicted cells. The four same-section STARmap validation datasets include old and young tongue samples containing 2,217 and 2,641 cells, respectively, and old and young kidney samples containing 14,541 and 15,616 cells, respectively.

For the GenVinci-3DST evaluation, the generated aging human skin atlas consisted of 9 samples producing 13.2 million predicted cells. The four adjacent-section STARmap validation datasets include three sections from a young donor (23 years old) containing 8,133, 4,702, and 4,438 cells, respectively, and one section from an older donor (71 years old) containing 3,835 cells.

For the GenVinci-gsMap evaluation, we analyzed the same reconstructed human skin atlas data as in the GenVinci-3DST part. We obtained publicly available GWAS summary statistics for five skin diseases (AD, PsO, LP, BCC, cSCC) and two negative-control traits (MCV, SCZ; Table S3).

### Benchmarks for downstream tasks

For three classical transcriptomics tasks, including cell type annotation, spatial domain identification, and batch integration, we compared GenVinci with four state-of-the-art single-cell and spatial foundation models: Geneformer^17^ (scRNA-seq), scGPT^16^ (scRNA-seq), Nicheformer^19^ (spatial + scRNA-seq), and scGPT-spatial^18^ (spatial + scRNA-seq). All evaluations were performed in a zero-shot setting by extracting pretrained embeddings without model fine-tuning. For cell type annotation, we trained a linear probe using a logistic regression classifier with default parameters on the extracted embeddings.

For age prediction, we benchmarked GenVinci (using linear probing) against two baselines: (1) normalized gene expression without spatial smoothing (Single-cell, SC) and (2) SpatialAgingClock^23^ (SAC), the current state-of-the-art method for age prediction on spatial transcriptomics data. SpatialAgingClock employs SpatialSmooth, which constructs local cell graphs and iteratively smooths gene expression across spatially neighboring cells before age regression, and its authors demonstrated that such spatial smoothing substantially improves age prediction performance.

To assess the contribution of spatial context, we evaluated two variants of our method: SC + GenVinci, which uses embeddings derived directly from normalized gene expression, and SAC + GenVinci, which incorporates SpatialSmooth-based neighborhood aggregation on the embeddings. We evaluated both methods under two settings: (1) cell-type-specific leave-one-mouse-out cross-validation on the coronal dataset and (2) cross-dataset generalization by training on the coronal dataset and testing on the independent sagittal dataset. All methods used a Lasso regression for age prediction.

For senescence identification, we compared GenVinci against seven representative benchmark approaches spanning single-marker, gene-set, and machine learning methods. The single-marker strategy uses log-normalized expression values of canonical senescence markers such as p16 and p21, assigning higher scores to cells with elevated expression. We used GenVinci to extract gene expression embeddings and train a linear classifier to predict the held-out samples. The gene-set ranking approach leverages published senescence gene sets such as SenMayo^26^ and hUSI^25^, applying enrichment methods including ssGSEA (GSVA R package) and AUCell to calculate continuous senescence scores. The SenCID^28^ framework employs a machine-learning pipeline to derive both continuous and binary senescence scores directly from expression count matrices, offering improved generalization over fixed gene signatures. DeepScence^27^ uses an autoencoder-based architecture trained on large curated senescence gene sets. HVG-PCA provides a strong, matched supervised baseline by training a logistic regression classifier on principal components derived from highly variable genes using the same labels. These benchmarks represent diverse strategies, from simple marker-based scoring to advanced machine learning, enabling comprehensive evaluation of GenVinci’s ability to capture senescence and aging signals.

For senescence classifier and SenNet validation, we mainly benchmarked with SenMayo and canonical senescence markers (p16, p21) or experiment-defined senescence labels. The scores are computed using *the scanpy.tl.score_genes* function.

For gene expression prediction from histology, we compared GenVinci against several representative methods spanning local, spatial, contrastive, and baseline strategies. ST-Net^13^ (local) fine-tuned pretrained convolutional networks (DenseNet-121) on local image patches to predict spot-level expression but lacks global context. Hist2ST^21^ (spatial) improves upon this by incorporating spatial graphs to model neighborhood relationships between patches, while HisToGene^20^ (spatial) emphasizes spatial consistency by jointly optimizing patch-level features with global tissue architecture via a vision-transformer architecture. BLEEP^22^ takes a different approach by using contrastive learning to align histology and expression modalities in a shared latent space, enhancing cross-sample generalization. HEST^32^ provides a minimal baseline of our model that directly maps the same pretrained pathology foundation model embeddings to expression values using a ridge regression with PCA (256 factors) without explicit spatial or alignment components. These benchmarks represent diverse methodological paradigms, allowing rigorous evaluation of GenVinci’s advantages.

For gsMap-based analyses, we compared our approach with the original gsMap implementation and evaluated the contributions of incorporating GenVinci embeddings and our improved continuous, magnitude-aware marker scoring scheme designed to support dense transcriptomic matrices.

### Evaluation metrics

For cell type annotation, we reported accuracy, weighted F1 score, area under the receiver operating characteristic curve (AUROC), and confusion matrices. For each dataset, cells were randomly split into 80% for training and 20% for held-out testing.

For spatial domain identification, we benchmarked unsupervised embeddings against manual annotations using the adjusted Rand index (ARI), which measures label concordance while adjusting for random chance, and the normalized mutual information (NMI), which captures shared information between predicted and true cluster assignments.

For batch integration, we adopted the scIB benchmarking framework^85^ following previous studies and reported NMI, ARI, ASW_label, AvgBIO, and AvgBATCH. AvgBIO represents the average of multiple biological conservation metrics, including NMI, ARI, and silhouette scores, and measures how well the embeddings preserve underlying biological structure. AvgBATCH represents the average of batch correction metrics, including ASW and graph connectivity, and evaluates how effectively embeddings integrate batches while minimizing technical confounders.

For age prediction, we evaluated at both cell and mouse levels. For mouse-level evaluation, predicted ages were averaged across cells from the same mouse, and we additionally employed a cell-type ensemble strategy by first averaging predictions within each cell type and then across cell types to reduce biases arising from differences in cellular composition. Performance was assessed using mean absolute error (MAE) and coefficient of determination (*R*²).

Senescence identification performance was mainly evaluated using the area under the receiver operating characteristic curve (AUROC), because cellular senescence is a continuous phenotype and there is no universal threshold to define binary senescent and non-senescent states across all the benchmarks.

Model performance for spatial gene expression prediction was measured using the average Pearson correlation coefficient (PCC) between predicted and ground-truth expression profiles within each dataset. We also used ARI and NMI to measure the clustering performance. For human skin samples with adjacent STARmap sections, we assessed the biological fidelity of H&E-predicted virtual transcriptomics by comparing neighboring tissue sections from the same specimen in the absence of paired ground-truth measurements. Agreement was evaluated at the gene and cell-type levels using both distribution-based metrics (total variation distance and Jensen–Shannon distance) and correlation-based metrics (Pearson and Spearman correlations). Lower distances and higher correlations indicate improved reconstruction fidelity and cross-section reproducibility. For virtual section generation, we computed the same distance- and correlation-based metrics.

For comparison with same-section STARmap measurements in mouse tongue and kidney (Figure 4E and Figure S6D–H), we summed all STARmap transcripts inside a 100 × 100 µm window centered on each cell, then normalized and log-transformed the counts. We predicted expression for the same cells from H&E patches centered on each cell. Prediction and measurement were each smoothed by averaging over neighboring cells inside the same 100-µm window. Per-gene Pearson correlation was computed across cells for the 50 most highly variable genes (HVGs) in the measured data. For domain comparison, we clustered the predicted and measured profiles separately with the same pipeline (scaling, 50 principal components, 15-nearest-neighbor graph, Leiden resolution 0.5). The number of clusters k was not fixed in advance but was determined by the fixed resolution. Agreement was scored with ARI and NMI; because both metrics are invariant to label permutation, predicted domains did not need to be matched to measured ones.

For GWAS–spatial association mapping, we followed the gsMap framework and computed gene specificity scores (GSS) for each cell. Trait associations were quantified using stratified LD score regression (S-LDSC)^83^, which yields a *z*-score per cell for each trait. Because absolute significance was not calibrated on dense inputs, cell-type associations were summarized by fold enrichment: the fraction of cells in a cell type with *z* > 1.645 (one-sided *P* < 0.05) divided by the fraction among all other cells in the same section, with a pseudocount of one. An association was considered supported when fold enrichment exceeded 1.5, at least five cells in the cell type exceeded the threshold, and the enrichment exceeded that of both negative-control traits (mean corpuscular volume and schizophrenia) in the same cell type. The fold-enrichment cutoff excludes cell types only marginally above the section-wide background; the numbers of supported associations were essentially unchanged for cutoffs between 1.5 and 2. Seven literature-supported associations were evaluated (Results). Enrichment magnitudes were not compared across inputs or traits, and color scales are independent per panel.

### Implementation details

The GenVinci foundation model was pretrained using a 12-layer transformer encoder with a hidden dimension of 512, 8 attention heads, a feed-forward dimension of 2,048, and a maximum sequence length of 1,024. The model was trained with a batch size of 64 using AdamW optimization, a learning rate of 5 × 10^⁻⁵^, linear warmup for 20,000 steps, dropout of 0.10, and bf16 mixed precision. Gene tokens and expression tokens were randomly masked with probabilities of 0.15 and 0.40, respectively.

The GenVinci-2DST model used a hidden dimension of 512 and was trained with AdamW using a learning rate of 1 × 10^⁻⁴^, weight decay of 1 × 10^⁻⁴^, batch size of 256, and 30 training epochs. Model performance was monitored on validation datasets using valid-gene MSE and embedding cosine similarity.

## Data availability

For representation learning benchmarks, we used the mouse brain MERFISH dataset for cell type annotation (https://cellxgene.cziscience.com/collections/0cca8620-8dee-45d0-aef5-23f032a5cf09), the 10x Genomics human brain Visium dataset (https://github.com/lieberinstitute/humanpilot) for spatial domain identification, and the developing human thalamus MERFISH dataset (https://datadryad.org/dataset/doi:10.5061/dryad.98sf7m0q1) for batch integration. For age and senescence analysis, we used the mouse brain aging MERFISH datasets (coronal and sagittal) (https://zenodo.org/records/13883177) for age prediction, the 10x Visium mouse muscle injury dataset generated by notexin injection (https://datadryad.org/stash/dataset/doi:10.5061/dryad.t4b8gtj34), and the human brain Stereo-seq dataset (GSE269906; https://www.ncbi.nlm.nih.gov/geo/query/acc.cgi?acc=GSE269906) for senescence identification. For senescence validation, the human Drop-seq dataset is from (GSE102090; https://www.ncbi.nlm.nih.gov/geo/query/acc.cgi?acc=GSE102090), the WI-38 human fibroblast dataset is from (GSE226225; https://www.ncbi.nlm.nih.gov/geo/query/acc.cgi?acc=GSE226225), the murine lung tumor dataset is from (GSE203447; https://www.ncbi.nlm.nih.gov/geo/query/acc.cgi?acc=GSE203447), and the murine brain microglia dataset is from (GSE229553; https://www.ncbi.nlm.nih.gov/geo/query/acc.cgi?acc=GSE229553). For GWAS enrichment analysis, we collected publicly available summary statistics from the GWAS Catalog (https://www.ebi.ac.uk/gwas/downloads/summary-statistics), FinnGen R12, Pan-UKB, and the Psychiatric Genomics Consortium (Table S3).

For model pretraining, we curated publicly available single-cell and spatial transcriptomics datasets from 10x Genomics (https://www.10xgenomics.com/datasets), CELLxGENE (https://cellxgene.cziscience.com/datasets), HEST-1k (https://huggingface.co/datasets/MahmoodLab/hest), SODB (https://gene.ai.tencent.com/SpatialOmics/), STimage-1K4M (https://huggingface.co/datasets/jiawennnn/STimage-1K4M), SenNet (https://data.sennetconsortium.org/), HCA/HTAN (HTAPP) (https://data.humantumoratlas.org/explore), HuBMAP (https://portal.hubmapconsortium.org/search/datasets), NanoString CosMx (https://nanostring.com/products/cosmx-spatial-molecular-imager/ffpe-dataset/), Vizgen (https://vizgen.com/data-release-program/), and Allen Brain Map (https://knowledge.brain-map.org/data?limit=50&search=%2A&sort=default~ASC). Detailed data description and manually curated dataset information can be found in Table S1.

## Code availability

The source code and Python package are available at https://github.com/jian-shu-lab/GenVinci.

## Author contributions

J.S. conceived, led, and supervised the project. Under the supervision of J.S., X.C. designed and developed the model, performed the benchmarking and downstream analyses, and wrote the manuscript. C.C. contributed to model development and scientific discussion. Y.W. generated and processed the human skin H&E imaging and STARmap data. Y.-L.S. generated and processed the mouse atlas H&E imaging and STARmap data. J.Zhou contributed to figure preparation and manuscript editing. H.H. and F.M. contributed to sequencing data analysis and manuscript editing. S.Sapantzi, M.C., J.Q., X.L., Y.C., Y.Z., and J.Ziegler reviewed the manuscript and contributed to scientific discussion. R.T. performed H&E slide imaging and scanning. K.T. and S.Sol provided the human skin samples. S.D., A.M., W.M., D.S., C.L., J.C.S., H.P., Z.D., K.Z., J.W.K., H.Y., A.R., C.M.S., and P.T.C.S. reviewed the manuscript and contributed to scientific discussion. All authors reviewed and approved the manuscript.

## Acknowledgements

The data analyses of this work were partially conducted on the MLSC Compute Cluster and ERISXdl Linux GPU Platform at Massachusetts General Hospital, and the FASRC Cannon cluster supported by the FAS Division of Science Research Computing Group at Harvard University. This work was supported by NIH New Innovator Award (DP2TR004354), the Biswas Family Foundation, Massachusetts Life Sciences Center, and Massachusetts General Hospital to J.S., NIH Common Fund (UG3CA275687, UH3CA275687) to J.S., P.T.C.S., J.W.K., MIT Generative AI Impact Consortium to J.S., P.T.C.S. We thank the NIH Common Fund Cellular Senescence Network (SenNet) Consortium and its tissue mapping centers and technology development sites for generating and sharing the data accessed through the SenNet Data Portal. We thank Brendan Honick at the Pittsburgh Supercomputing Center, Carnegie Mellon University, for help in navigating and downloading the SenNet data.

## Declaration of generative AI and AI-assisted technologies in the writing process

During the preparation of this work, the authors used ChatGPT and Claude in order to improve language. After using these tools or services, the authors reviewed and edited the content as needed and take full responsibility for the content of the publication.

---

**Figure S1.**
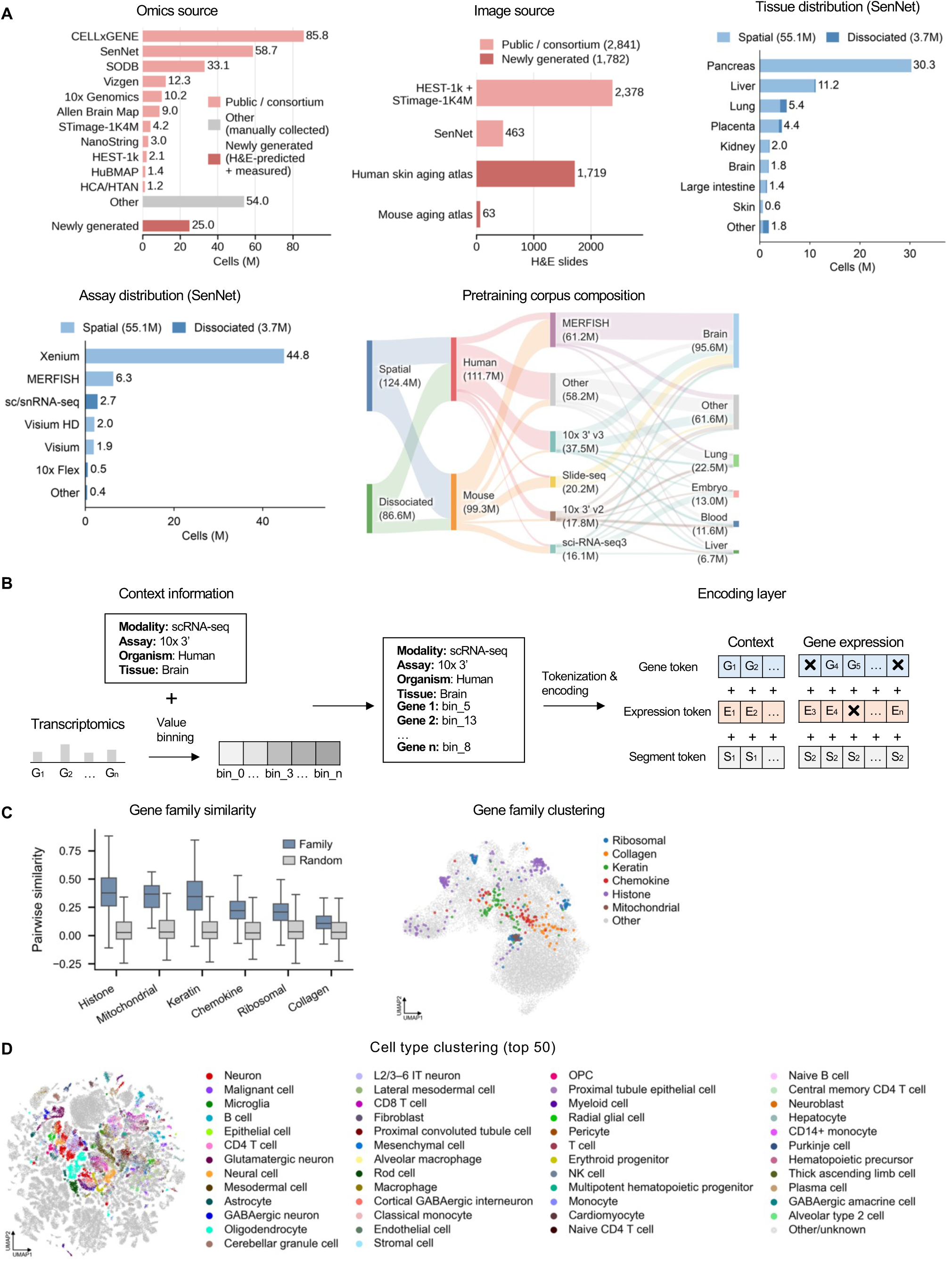
Composition of ImageOmicsNet-300M and characterization of GenVinci representations. **A,** Composition of ImageOmicsNet-300M. Omics source, number of transcriptomic profiles (millions) contributed by each data source: public and consortium resources (pink), other manually collected datasets from publications and consortia (gray), and newly generated data, comprising H&E-predicted virtual spatial profiles and measured STARmap and snRNA-seq data (red). Image source, number of H&E sections from public and consortium resources (2,841) and from the newly generated mouse and human skin aging atlases (1,782). Tissue distribution (SenNet) and assay distribution (SenNet), number of cells (millions) in the October 2026 SenNet data release by tissue and by assay, colored by spatial (55.1 million) and dissociated (3.7 million) profiles. Pretraining corpus composition, Sankey diagram of the 211.0 million experimentally measured profiles used for foundation-model pretraining, by modality, species, assay, and tissue; HEST-1k, STimage-1K4M, and the October 2026 SenNet release were excluded from pretraining, and the earlier SenNet release (0.8 million cells) was included. Values are rounded to one decimal place and may not sum exactly to the totals shown. **B,** Encoding strategy used for GenVinci pretraining. Biological context, including modality, assay, organism, and tissue, is combined with binned transcriptomic profiles to generate context, gene, expression, and segment tokens. **C,** Characterization of learned gene embeddings. Left, distributions of pairwise cosine similarity within six gene families, compared with size-matched random gene sets excluding family members (Random: 20 replicates per family, up to 5,000 pairs per box). Families are ordered by median difference. Boxes show median and interquartile range; whiskers, 1.5× interquartile range; outliers omitted. Right, UMAP of the gene embedding space highlighting the same gene families. **D,** UMAP of learned cellular representations colored by the 50 most abundant annotated cell types. Cells are the same 100,000-cell subsample as in Figure 1C; annotations are as provided by the source datasets, without harmonization.

**Figure S2.**
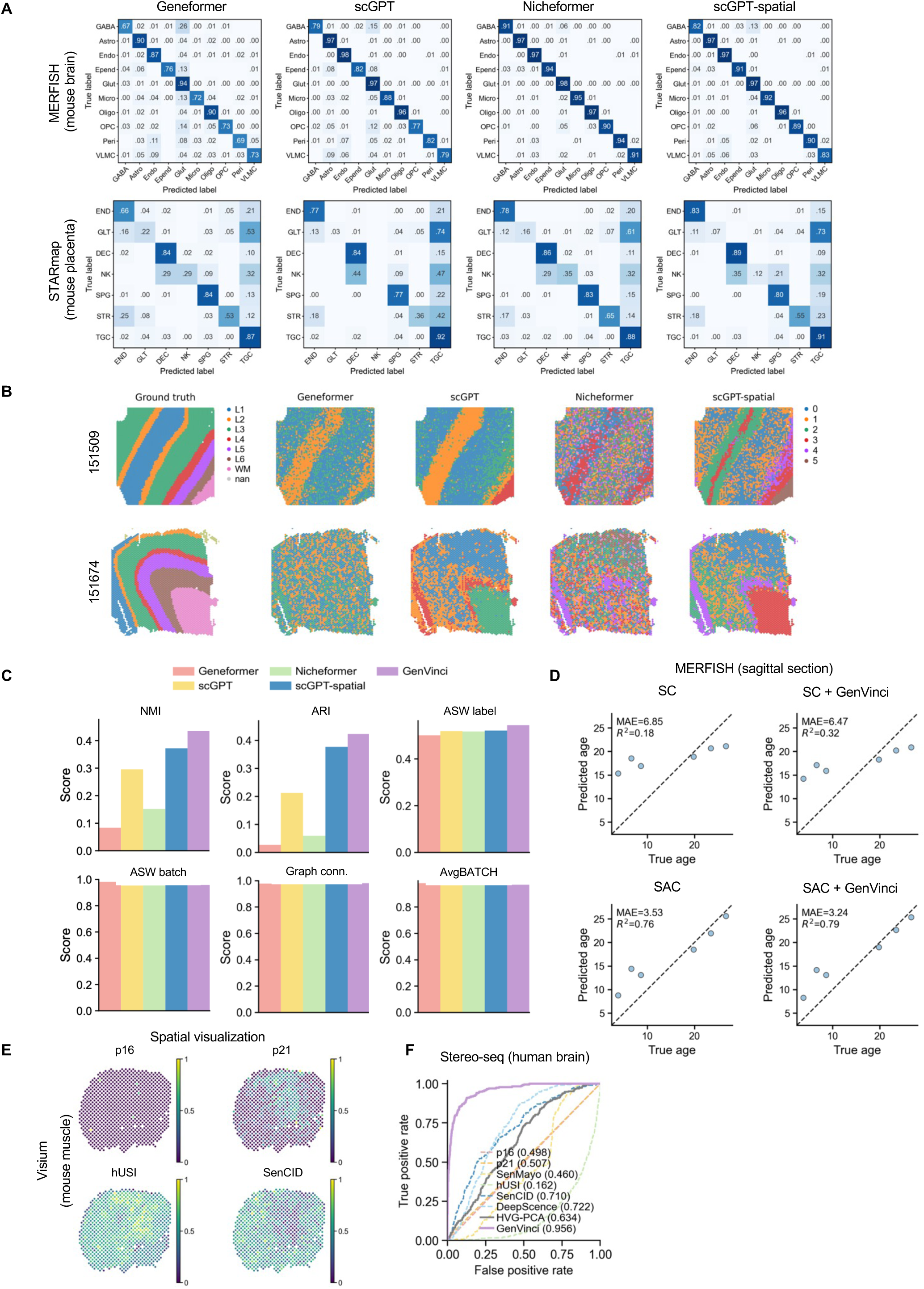
Additional benchmarking of GenVinci across downstream tasks. **A,** Confusion matrices for cell-type annotation on MERFISH mouse brain and STARmap mouse placenta datasets across four competing methods. Abbreviations as in Figure 2A. **B,** Spatial domain identification on two Visium human brain samples by four competing methods. **C,** Quantitative comparison of batch integration performance on the MERFISH human developmental thalamus dataset, measured by NMI, ARI, and ASW label (top: biological conservation) and ASW batch, graph connectivity, and AvgBATCH (bottom: batch correction). ASW, average silhouette width. **D,** Age prediction on the independent MERFISH sagittal mouse brain dataset. Models were trained on the coronal dataset. **E,** Spatial distributions of predicted senescence scores on the mouse muscle Visium dataset; the remaining methods are shown in Figure 2E. **F,** ROC curves comparing competing methods on the human brain Stereo-seq dataset, plotted as in Figure 2E.

**Figure S3.**
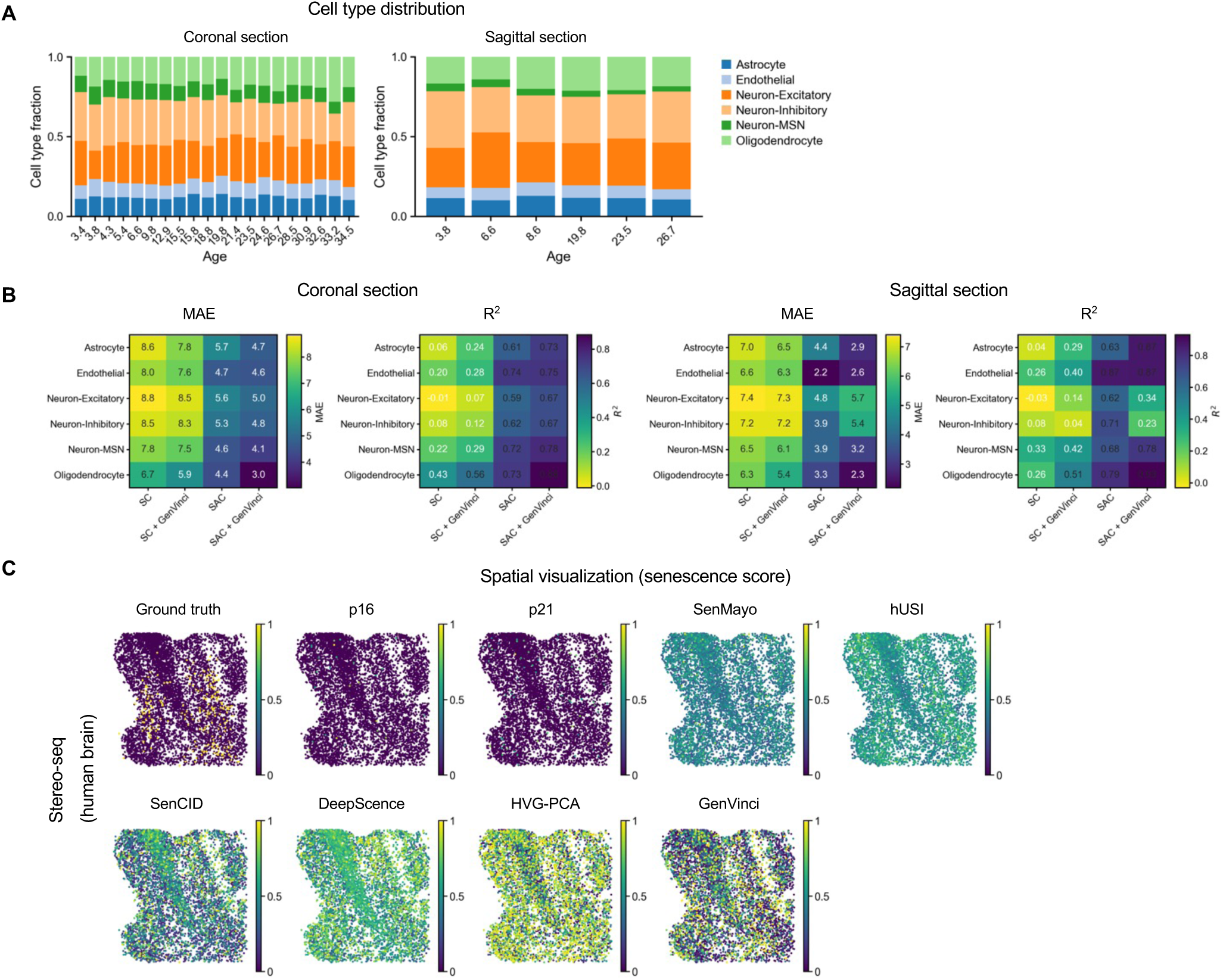
Additional evaluation of age prediction and senescence identification. **A,** Cell-type composition across chronological ages in the MERFISH mouse brain datasets used for age prediction. **B,** Cell-type-specific age prediction performance. Heatmaps show the mean MAE and *R*² for SC, SAC, and the corresponding GenVinci-based variants across six major cell types in the coronal (left) and sagittal (right) sections. Color scales are set independently for each panel. **C,** Spatial distributions of predicted senescence scores in the human brain Stereo-seq dataset.

**Figure S4.**
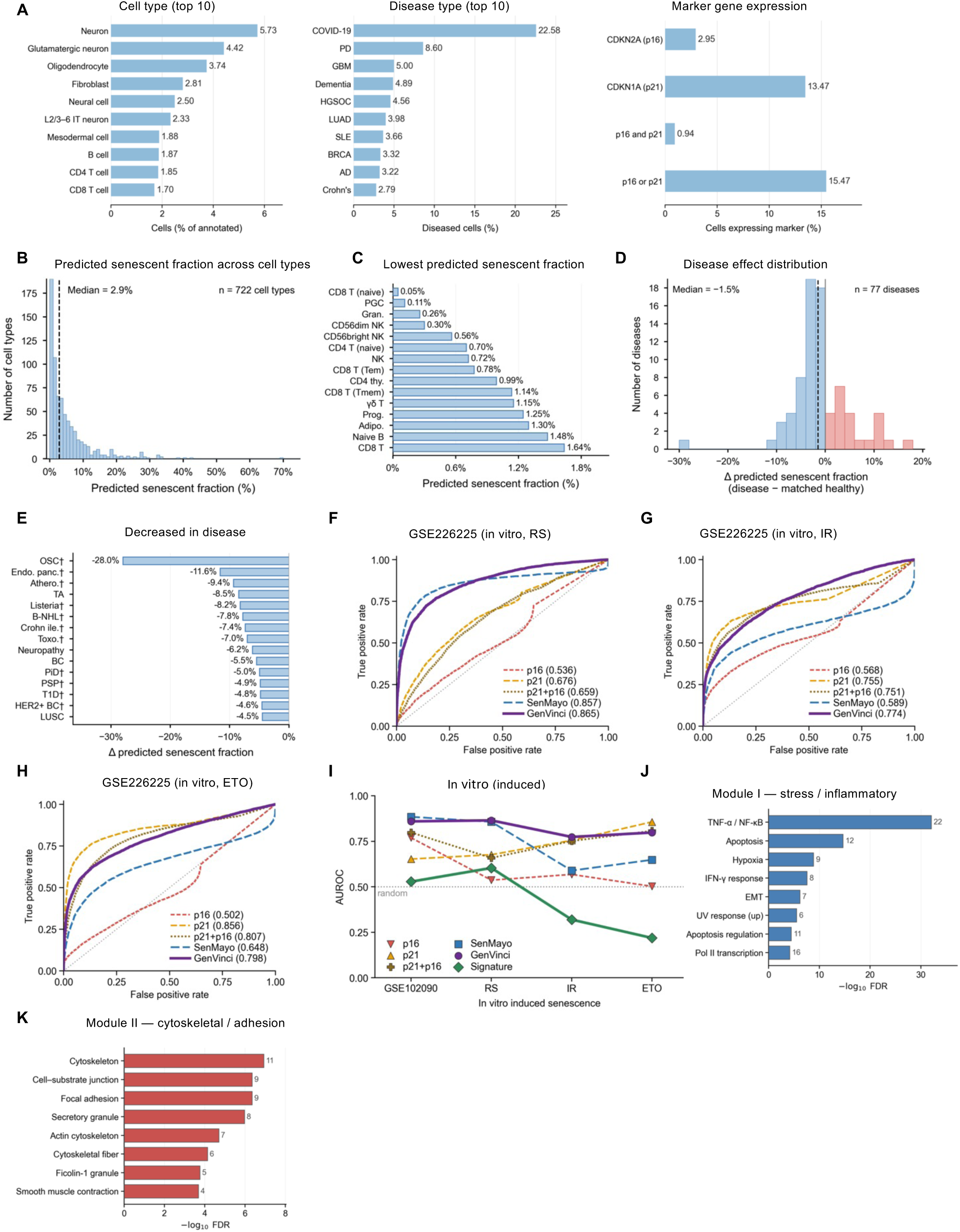
Additional characterization of the atlas-wide senescence projection and its external validation. **A,** Composition of the atlas used for the senescence projection. Left, the ten most abundant annotated cell types. Middle, the ten most abundant disease states. Right, the proportion of cells in which *CDKN2A* (p16), *CDKN1A* (p21), both, or either marker is detected. **B,** Distribution of predicted senescent fraction (proportion of cells in the top 5% of S; Figure 3A) across all 722 cell types with at least one qualifying healthy group (≥1,000 cells). **C,** The 15 cell types with the lowest predicted senescent fraction, plotted as in Figure 3B. **D,** Distribution of disease effects on predicted senescent fraction across 77 diseases, computed as in Figure 3D. Bars are colored by the direction of the effect. **E,** The 15 diseases with the largest decrease in predicted senescent fraction relative to matched healthy samples, plotted as in Figure 3D. **F–H,** ROC curves for the identification of senescent cells in three *in vitro* induction arms of the GSE226225 dataset: replicative senescence (F: RS vs. CTRL), irradiation-induced senescence (G: IR vs. CTRL), and etoposide-induced senescence (H: ETO vs. CTRL), plotted as in Figure 3F. **I,** AUROC across all four *in vitro* comparisons for marker-based detection (p16, p21, and their union), gene-set scoring (SenMayo), the senescence classifier (GenVinci), and SenNet-100. The dotted line indicates random performance. **J, K,** Over-representation analysis of the two principal co-variation modules identified in Figure 3G: Module I (J) and Module II (K). Libraries: GO Biological Process 2023, GO Cellular Component 2023, Reactome 2022, and MSigDB Hallmark 2020. Cell-type and disease abbreviations as in Figure 3, with the following additions—PGC, primordial germ cell; Gran., granulocyte; Prog., progenitor cell; Adipo., adipocyte; CD4 thy., CD4-positive, alpha-beta thymocyte; Tem, effector memory T cell; Tmem, memory T cell; OSC, malignant ovarian serous tumor; Endo. panc., endocrine pancreas disorder; Athero., atherosclerosis; TA, tubular adenoma; *Listeria*, listeriosis; B-NHL, B-cell non-Hodgkin lymphoma; Crohn ile., Crohn ileitis; Toxo., toxoplasmosis; BC, breast cancer; PiD, Pick disease; PSP, progressive supranuclear palsy; T1D, type 1 diabetes mellitus; HER2+ BC, HER2-positive breast carcinoma.

**Figure S5.**
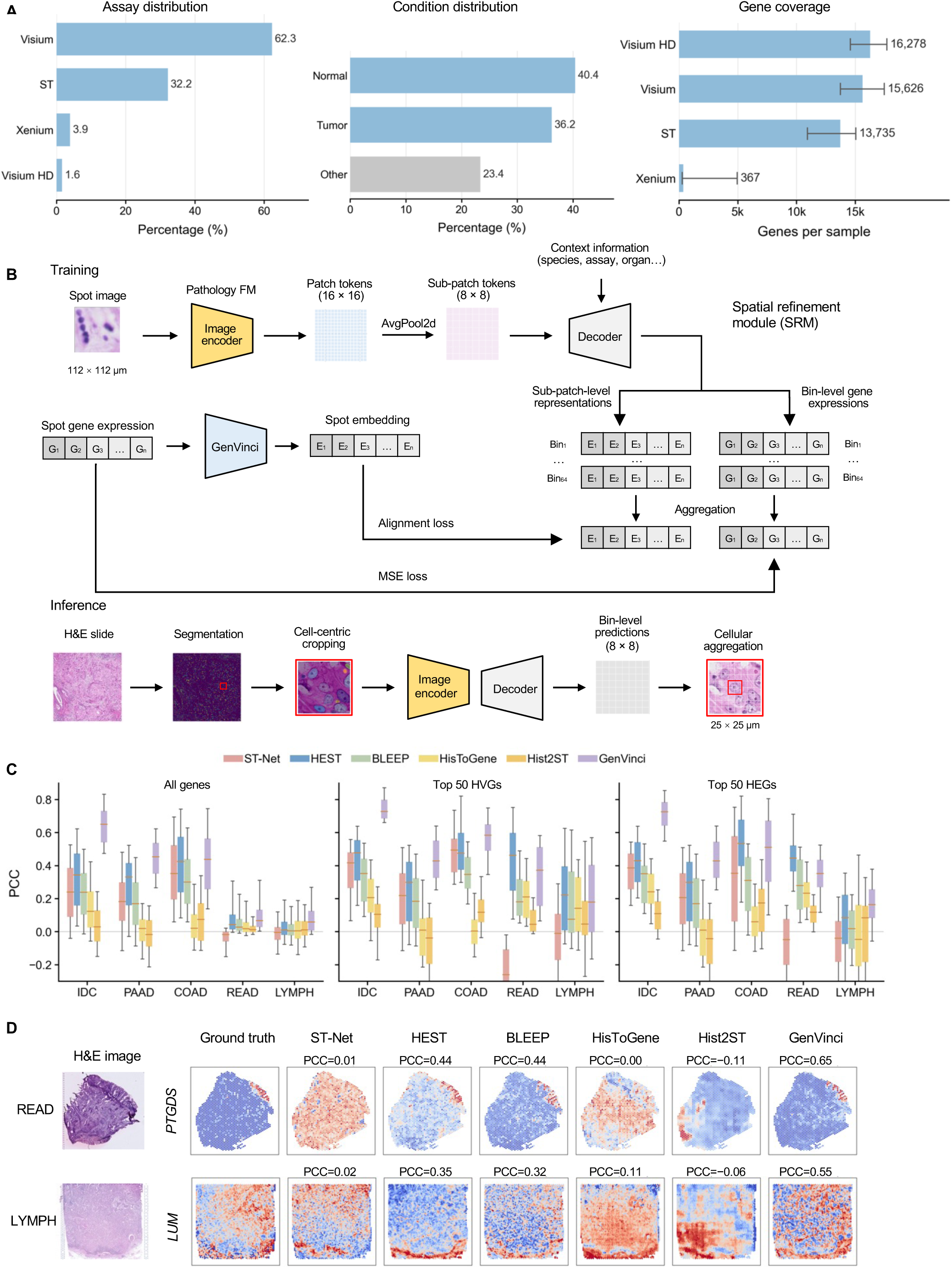
Architecture and out-of-distribution evaluation of GenVinci-2DST. **A,** Composition of the ImageOmicsNet data used to train GenVinci-2DST, showing distributions across assays, biological conditions, and gene coverage. **B,** Detailed architecture of GenVinci-2DST training and inference. During training, H&E patches are encoded by a pathology foundation model and decomposed into sub-patch-level representations by the Spatial Refinement Module (SRM), while GenVinci embeddings derived from matched spatial transcriptomics provide teacher supervision. During inference, cell-centric H&E crops are encoded and decoded into bin-level expression profiles, with averaging over the central 2 × 2 bins yielding cellular-level virtual transcriptomics. **C,** Distribution of gene-expression prediction performance across held-out IDC, PAAD, COAD, rectal adenocarcinoma (READ), and axillary lymph nodes from patients with IDC (LYMPH) datasets for all genes (left), the top 50 HVGs (middle), and the top 50 HEGs (right). **D,** Representative spatial gene-expression predictions for held-out READ and LYMPH samples.

**Figure S6.**
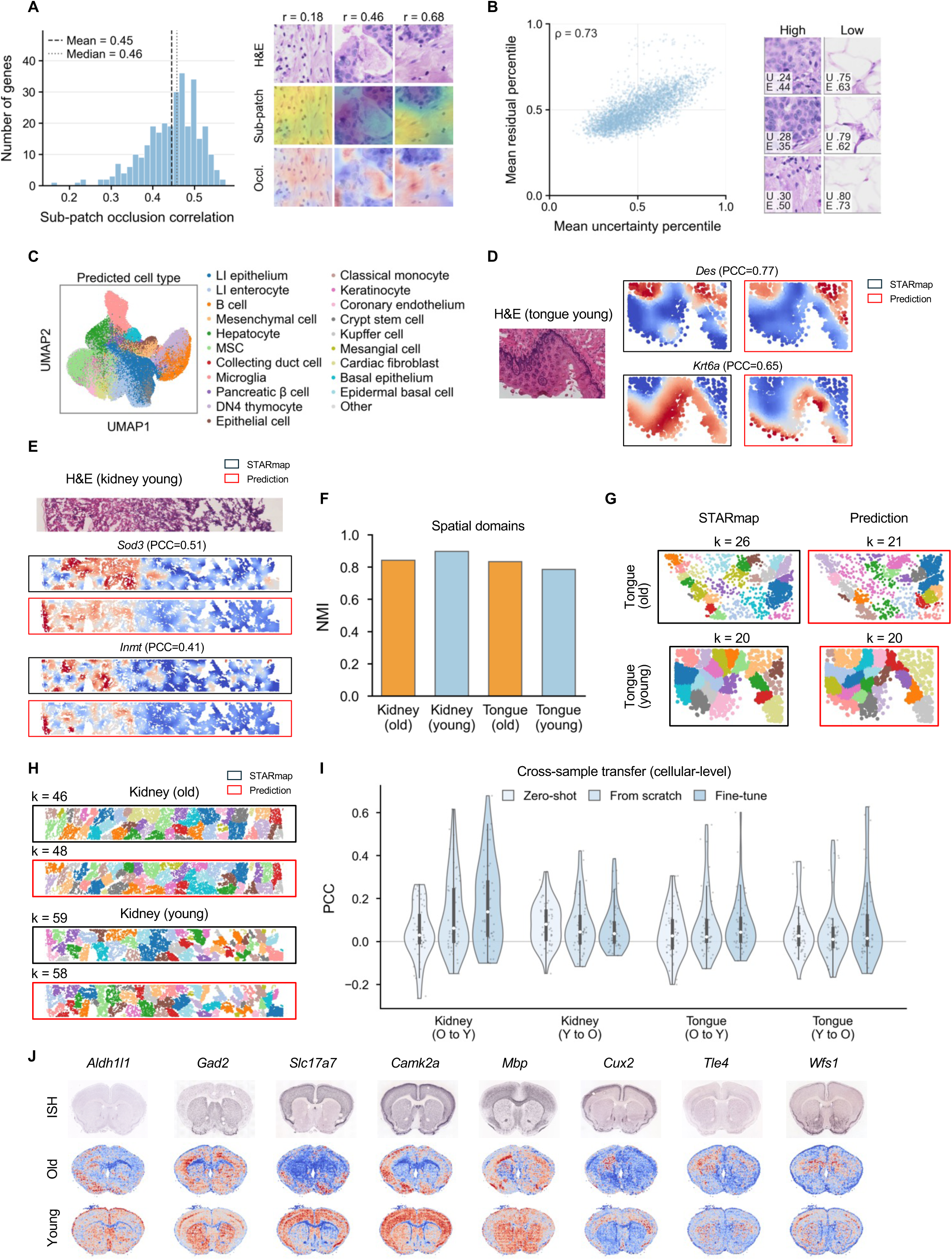
Model interpretability, prediction uncertainty, and validation of the virtual transcriptomic atlas. A, Leave-one-sub-patch-out occlusion analysis. Each 224 × 224 px (112 × 112 μm) H&E patch centered on a 100-μm spot was divided into an 8 × 8 grid of non-overlapping 28 × 28 px sub-patches, each corresponding to one expression bin, and each sub-patch was occluded in turn to measure its contribution to the spot-level prediction (prediction with all sub-patches minus prediction without sub-patch *k*). Left, distribution of the sub-patch–occlusion correlation across all genes; for each gene, the Pearson correlation between the 64 bin-level expression predictions and their corresponding occlusion effects was computed within each spot and summarized as the median across spots (dashed line, mean across genes; dotted line, median across genes). Right, representative spots for *POSTN* selected at the 10th, 50th, and 90th percentiles of the within-spot Pearson correlation (*r*). Rows show the H&E patch, the bin-level expression prediction map, and the occlusion effect map (red, positive; blue, negative), overlaid on the patch. **B,** Test-time augmentation (TTA) uncertainty. Uncertainty is the standard deviation of predictions across nine spatial shifts of each patch (0 and ±16 px along both axes). Left, spot-level relationship between uncertainty and absolute prediction error, each gene-wise rank-normalized to a percentile and averaged across genes. Right, representative patches drawn from the lowest (High confidence) and highest (Low confidence) deciles of mean uncertainty percentile. *U*, mean uncertainty percentile; *E*, mean residual percentile. *r*, Pearson correlation coefficient; ρ, Spearman correlation coefficient. **C,** UMAP visualization of reconstructed predicted cells colored by predicted cell type. **D, E,** Additional comparisons of STARmap measurements and GenVinci-2DST predictions for representative marker genes in young tongue (D) and young kidney (E). **F,** Agreement between STARmap- and prediction-derived clustering (Leiden clustering, resolution 0.5; Methods), quantified by the normalized mutual information (NMI). **G, H,** Comparison of spatial domains identified from STARmap measurements and GenVinci-2DST predictions in old and young tongue (G) and kidney (H). *Spatial domains were identified by Leiden clustering (resolution 0.5) without fixing the number of clusters; k denotes the number of Leiden clusters recovered.* **I,** Cross-sample evaluation of GenVinci-2DST at cellular-level (25-μm) resolution. Models were trained on one STARmap sample and evaluated on a held-out sample from the other age group, in both directions, for kidney and tongue. Zero-shot, pretrained GenVinci-2DST; From scratch, a neural network model trained from scratch based on the H&E CLS embedding; Fine-tune, pretrained GenVinci-2DST fine-tuned on the training sample. Aggregation and smoothing as in Figure 4E, using 25-μm windows. Violin plots show the distribution of gene-level PCCs across the same 50 HVGs defined on the STARmap profiles. **J,** Comparison of GenVinci-2DST predictions in old and young mouse brain with the Allen Brain Atlas *in situ* hybridization (ISH) reference patterns for representative neuronal, glial, and regionally expressed genes. Each gene is scaled independently within each section.

**Figure S7.**
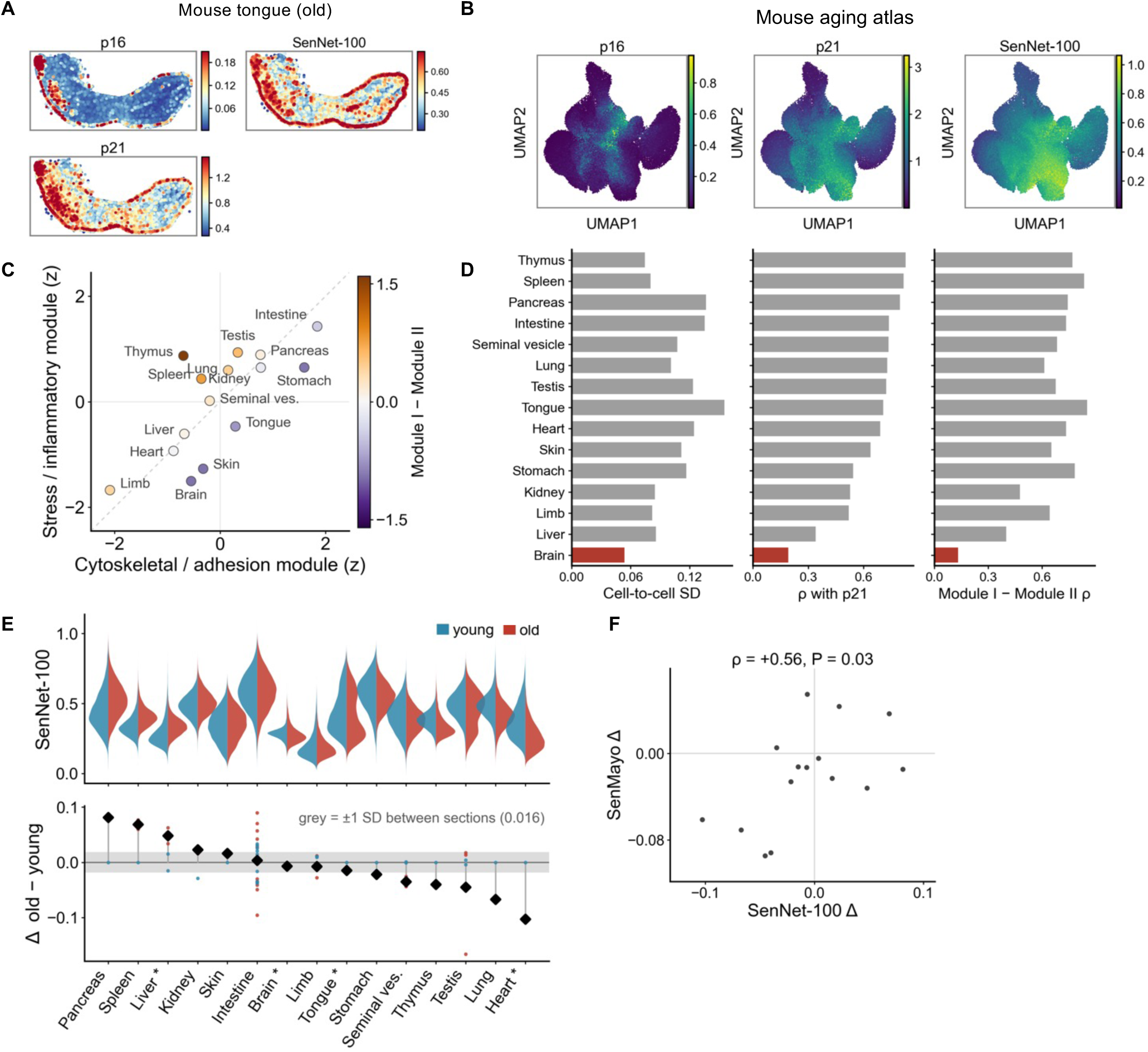
Senescence programs in the reconstructed aging mouse atlas. **A,** Spatial distribution of p16 (*Cdkn2a*), the SenNet-100 score, and p21 (*Cdkn1a*) in a representative old mouse tongue section. **B,** UMAP visualization of p16, p21, and the SenNet-100 score across the atlas. **C,** Dissociation of the two SenNet-100 modules across organs. **D,** Internal consistency of the SenNet-100 score per organ, quantified by cell-to-cell standard deviation, correlation with p21, and correlation between modules; brain is highlighted in red. **E,** Per-cell SenNet-100 distributions by organ, split by age (top) and per-organ difference in mean score between old and young mice (bottom); asterisks, organs not sampled from the main animal pair. **F,** Concordance of organ-level age differences estimated by SenNet-100 and SenMayo.

**Figure S8.**
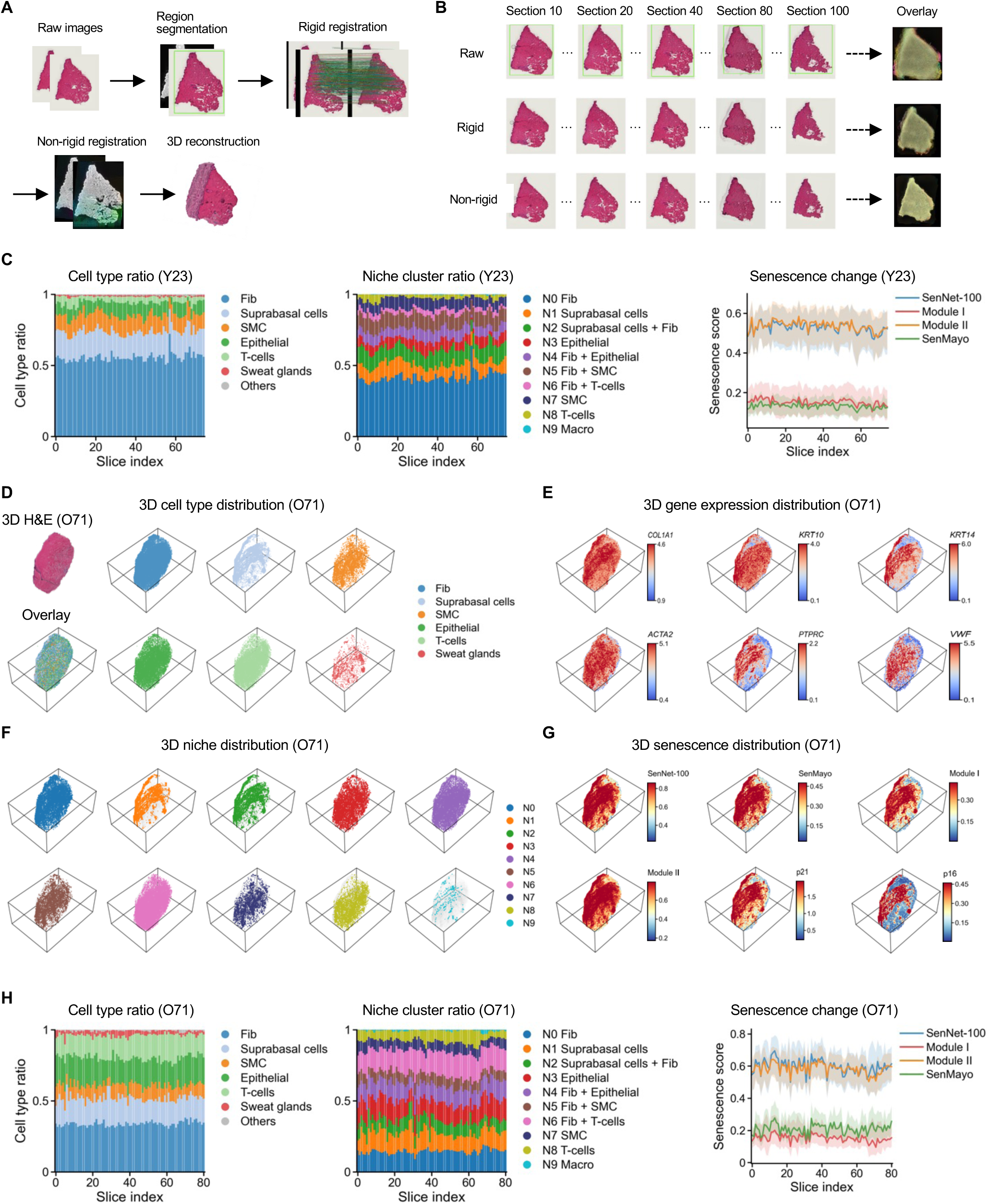
Three-dimensional reconstruction pipeline and per-section stability. **A,** Registration workflow: region segmentation, rigid alignment, non-rigid refinement, and three-dimensional reconstruction. **B,** Representative sections before registration (Raw), after rigid registration, and after non-rigid registration, with the corresponding overlays of all sections. **C,** Per-section composition for donor Y23 (23-1). Left, cell-type ratio; middle, niche ratio; right, senescence scores across sections (line, median; shading, interquartile range). **D–G,** As in Figure 5B–E, for donor O71 (71-1): three-dimensional H&E and cell-type distributions (D), marker gene expression (E), niches (F), and senescence distribution (G). **H,** Per-section cell-type ratio, niche ratio, and senescence scores for donor O71, as in C.

**Figure S9.**
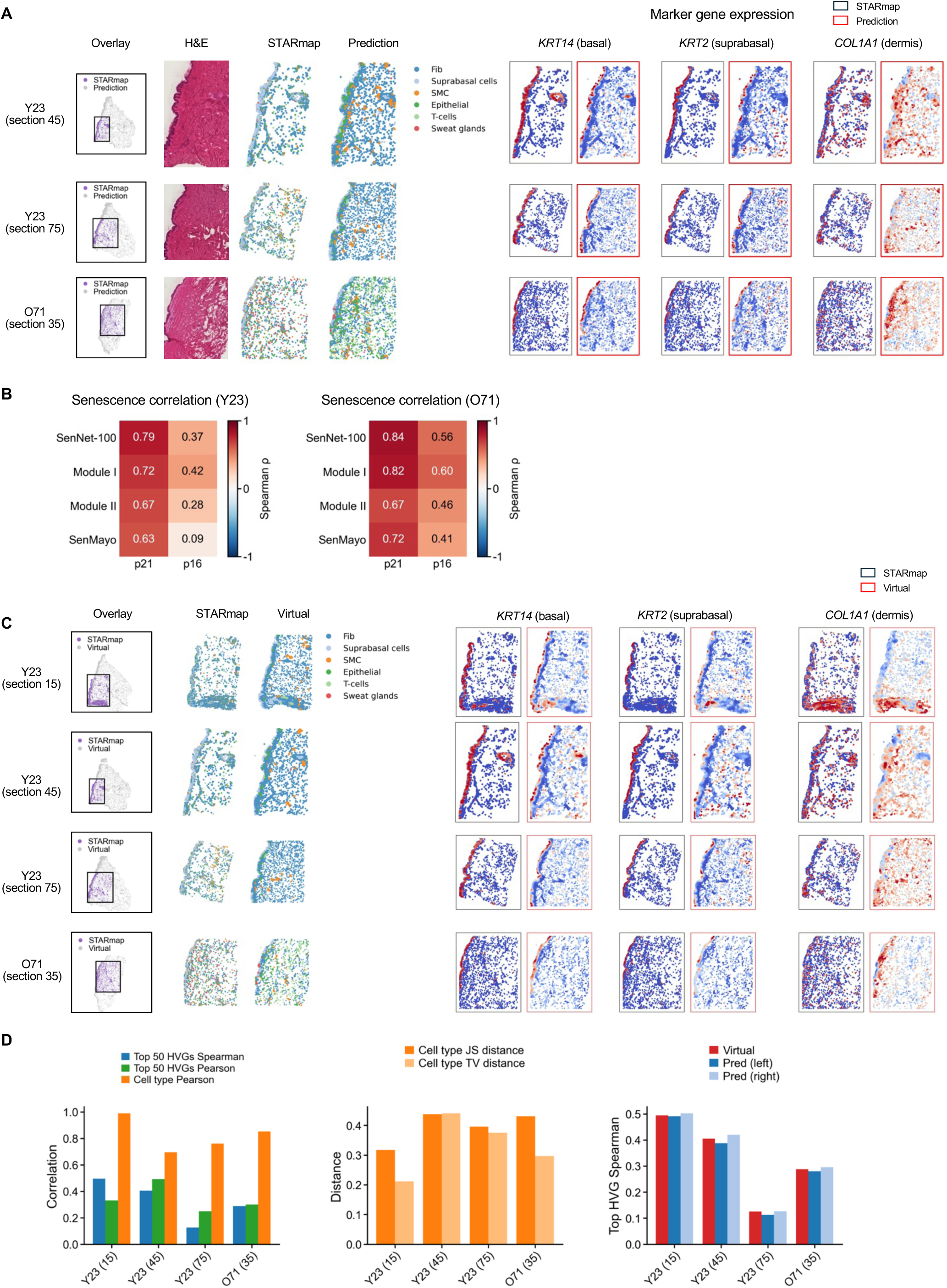
Validation against STARmap and evaluation of virtual section generation. **A,** Predicted versus STARmap sections for Y23 (sections 45 and 75) and O71 (section 35). For each: spatial overlap of the profiled region, H&E, cell-type maps, and paired expression of *KRT14* (basal), *KRT2* (suprabasal), and *COL1A1* (dermis). **B,** Spearman correlation between each senescence score (SenNet-100, Module I, Module II, and SenMayo) and p21 (*CDKN1A*) or p16 (*CDKN2A*) expression, computed across cells of donors Y23 and O71. **C,** Virtual sections generated at four held-out depths, compared with the corresponding STARmap sections, shown as in A. **D,** Quantitative agreement for the four sections. Left, correlation of the top 50 HVGs and of cell-type composition; middle, Jensen–Shannon and total-variation distances between cell-type compositions; right, top 50 HVG correlation for the virtual section compared with the two flanking predicted sections.

**Figure S10.**
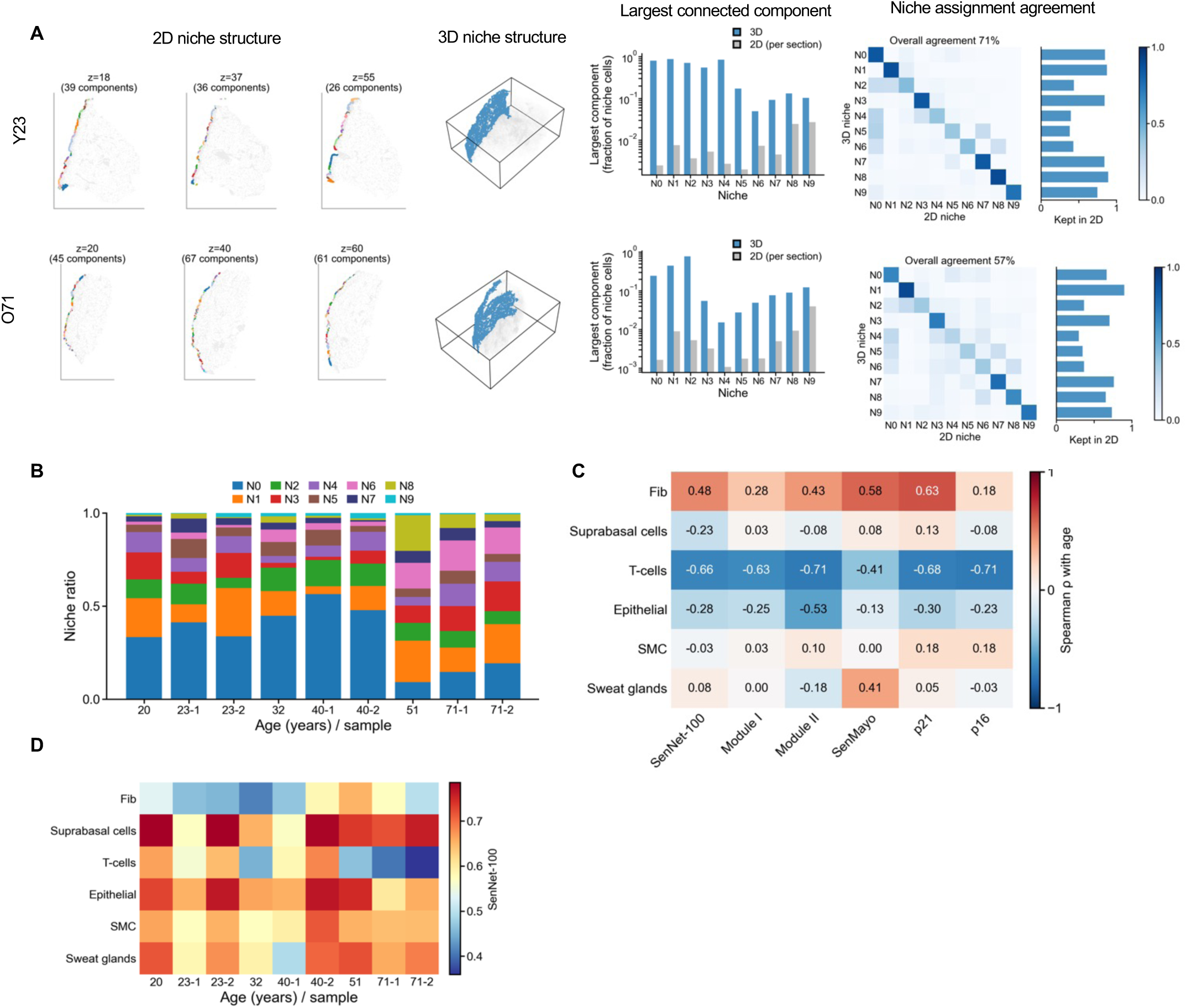
Three-dimensional reconstruction resolves structures fragmented in single sections. **A,** Niche structure in 2D versus 3D for donors Y23 (N1, epidermis) and O71 (N2, epidermal–dermal interface); connected components computed within a 30-μm radius. Left, cells of the largest 3D connected component shown in three sections, colored by within-section component (26–67 components per section). Middle, the same component in 3D (blue: 86.9% and 75.1% of niche cells), spanning all sections. Right, fraction of niche cells in the largest connected component in 3D versus single sections (log scale), and confusion matrix of niche assignment using 2D versus 3D neighborhoods (row-normalized; bars, fraction of cells retaining their label). **B,** Niche composition per donor, ordered by age. **C,** Spearman correlation between donor age and median senescence score per cell type (*n* = 9 donors). **D,** Median SenNet-100 score per cell type and donor, ordered by age.

**Figure S11.**
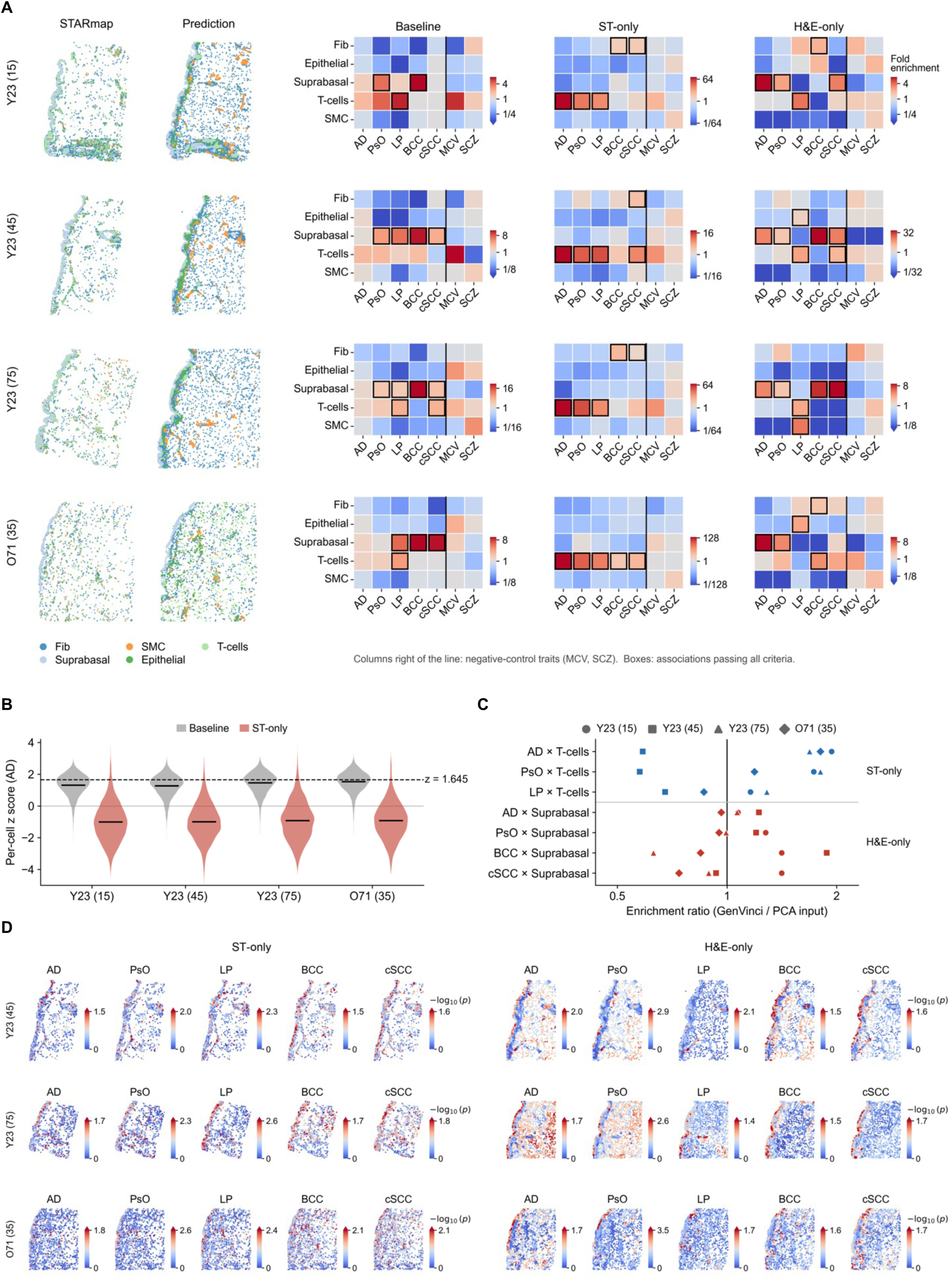
Validation of GenVinci-gsMap in sections with matched STARmap measurements. **A,** Spatial genetic mapping in the four sections with STARmap measurements, shown as in Figure 6B and including two negative-control traits (right of the vertical line). Black boxes, all associations meeting the criteria. **B,** Per-cell *z*-scores for AD with raw and imputed STARmap. Dashed line, *z* = 1.645. **C,** Ratio of enrichment obtained with GenVinci representations to that obtained with PCA as the graph input; each point is one section. **D,** Per-cell association maps for the three remaining sections, shown as in Figure 6C. Disease abbreviations as in Figure 6, with the following additions—MCV, mean corpuscular volume; SCZ, schizophrenia.

**Figure S12.**
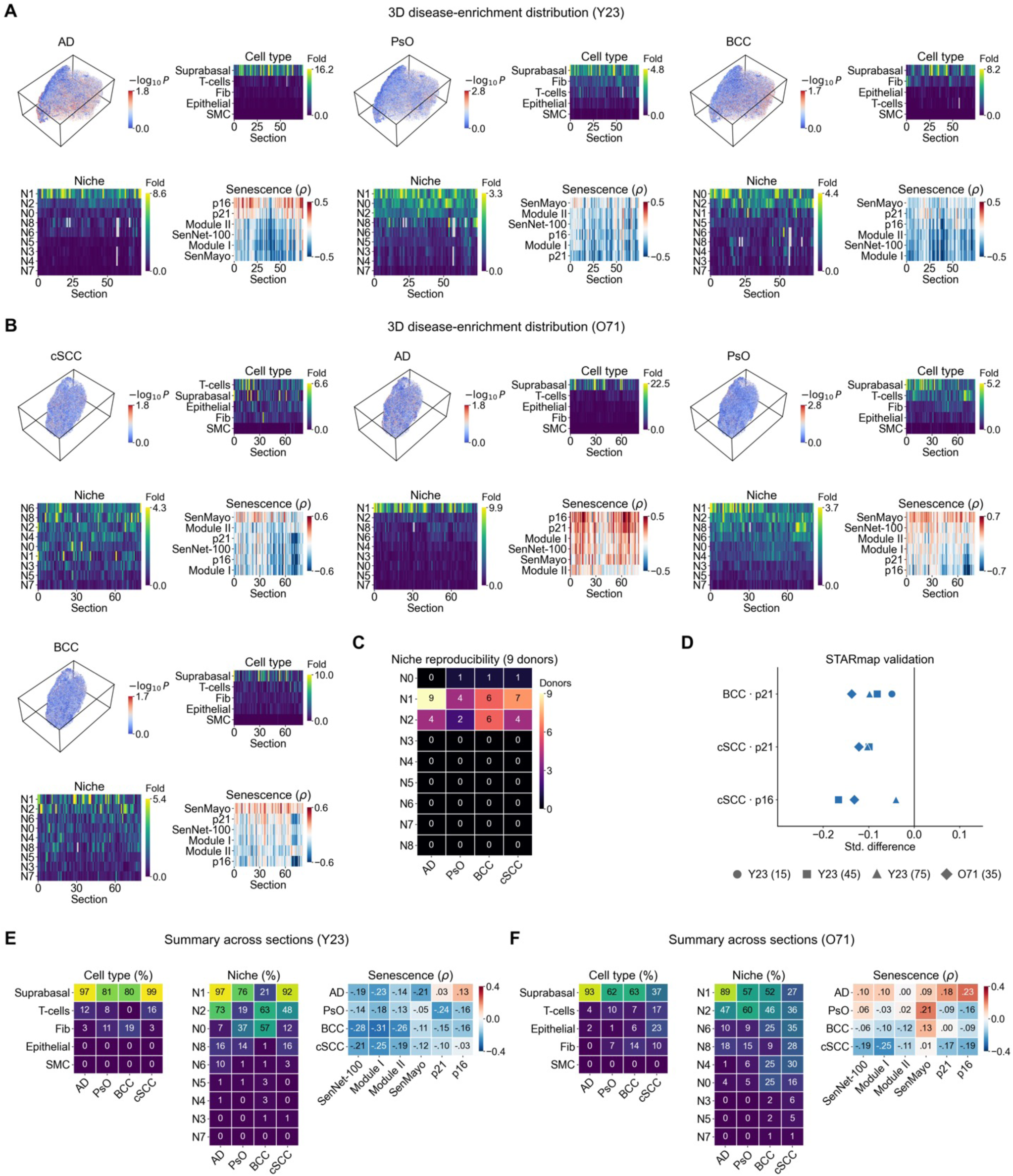
Three-dimensional disease-enrichment distributions within and across donors. **A,** Distributions for the remaining diseases in Y23, shown as in Figure 6E. **B,** Same as A for O71. **C,** Number of donors (of nine) in which each niche–disease association was supported. **D,** Difference in p21 or p16 expression between trait-associated and other suprabasal cells in the sections with STARmap measurements; each point is one section. **E, F,** Summaries across all sections of Y23 (E) and O71 (F): percentage of sections supporting each cell-type–disease and niche–disease association, and median correlation with each senescence measure. Niches as in Figure 6.

